# Synovial lining–expressed mechanosensor PIEZO1 drives inflammation-permissive macrophage phenotypes and joint inflammation

**DOI:** 10.64898/2026.01.12.699005

**Authors:** Theodoros Simakou, Marcus Doohan, Carmen Huesa, Kieran Woolcock, Domenico Somma, Watipenge Harmony Nyasulu, Shreya Prakash, Kieran Boyle, Luxin Yang, Binghua Darren Zhang, Denise Campobasso, Clara Di Mario, Andrew Farthing, Jack Frew, Lavinia Agra Coletto, Olympia Hardy, Lynette Dunning, Maria Rita Gigante, Luca Petricca, Viviana Antonella Pacucci, Scott Arkison, Yuriko Yamamura, Ivan De Martino, Maria Antonietta D’Agostino, Thomas D Otto, John Cole, Ruaidhri J Carmody, Carl Goodyear, Lucy MacDonald, Neil Basu, Stefano Alivernini, Mariola Kurowska-Stolarska

## Abstract

Synovial tissue–resident macrophages regulate immune homeostasis within the joint, but can adopt an inflammation-permissive phenotype that promotes immune cell infiltration in rheumatoid arthritis (RA). Understanding the factors that drive this phenotypic switch may help prevent the localisation of inflammation in the joints of individuals at-risk of RA. We identified the mechanosensitive ion channel PIEZO1 as a potential regulator of lining-layer synovial tissue macrophage (STM) function. PIEZO1 was highly expressed in homeostatic, tissue resident TREM2^pos^ lining-layer STMs and in its pathogenic chemokine producing TREM2^low^ phenotype that characterises the hyperplastic lining-layer in active RA. Intra-articular injection of a PIEZO1 agonist in mice induced neutrophil and monocyte infiltration, whereas inhibition of PIEZO1 signalling restored the protective macrophage phenotype. Thus, mechanosensing via PIEZO1 is a defining feature of the joint lining-layer, and its aberrant activation by mechanical stress may lead to the localisation of inflammation within the joint, facilitating a transition from asymptomatic autoimmunity of at-risk RA to clinical disease.

## Introduction

Rheumatoid arthritis (RA) begins with a breach of immune tolerance to citrullinated autoantigens in at-risk individuals, which can remain clinically silent (asymptomatic autoimmunity) for years before progressing to chronic joint inflammation and systemic comorbidities (Alivernini et al., 2022). Immune-targeted therapies have revolutionized the management of active disease, making remission achievable. However, most patients in remission still experience disease flares, resulting in significant pain and a progressive decline in quality of life (Alivernini et al., 2022). Therefore, strategies to prevent both the initial onset of disease and subsequent flares are urgently needed to reduce the personal and socio-economic burden of RA. Recent trials [TREAT_EARLIER (Krijbolder et al., 2022), APIPPRA (Cope et al., 2024) and PRAIRI (Gerlag et al., 2019)] in individuals at-risk-of RA have shown promise in delaying disease onset and reducing severity, but none have prevented its eventual development. These highlight the need for a deeper understanding of the mechanisms that drive the localisation of systemic asymptomatic autoimmunity specifically into the joints.

The synovial joint capsule is lined by the synovial membrane, a specialized tissue composed of two layers: the lining-layer, which faces the joint cavity, and the sublining-layer, which contains sensory neurons and blood vessels. Together, these layers provide immune protection and lubrication, enabling joint movement. In healthy synovium, resident lining-layer cells such as homeostatic TREM2^pos^VSIG4^pos^MERTK^pos^ (Cx3Cr1^pos^Trem2^pos^Vsig4^pos^Mertk^pos^ in mice) synovial tissue macrophages (STMs) and lubricin (PRG4)-producing CLIC5^pos^ fibroblast-like synoviocytes (FLS) form a protective, immunosuppressive barrier. In the sublining-layer, resident LYVE1^pos^ STMs establish a perivascular circuit with the THY1^pos^CXCL14^pos^GAS6^pos^ FLS cluster and pericytes, potentially regulating inflammatory cell extravasation and tissue repair (Alivernini et al., 2020a; Clayton et al., 2021; Croft et al., 2019; Culemann et al., 2019; Kurowska-Stolarska and Alivernini, 2022; MacDonald et al., 2024a; Schonfeldova et al., 2025).

In active RA, the synovial tissue undergoes pathological changes that promote the influx of inflammatory cells, leading to tissue hyperplasia and the formation of T/B cell-driven ectopic germinal centre–like structures (Alivernini et al., 2022). The synovial lining becomes hyperplastic, characterized by a decline in protective TREM2^pos^VSIG4^pos^MERTK^pos^ STM phenotypes, which are replaced by chemokine-producing pathogenic TREM2^low^SPP1^pos^ cells (Alivernini et al., 2020b; Bolton et al., 2025; Elmesmari et al., 2025). Simultaneously, the lining fibroblasts adopt a destructive phenotype, producing enzymes that degrade cartilage and bone (Croft et al., 2019). Experimental arthritis studies suggest that this shift in the lining-layer STM phenotype may trigger the localisation of inflammation in the joints, though the stimuli that drive this shift remain unknown (Schonfeldova et al., 2025; Zec et al., 2023a).

Synovial joints enable flexible and rapid motion, making them highly biomechanical organs, and emerging epidemiological evidence suggests that mechanical stress may contribute to the localisation of inflammation within the joints. Work-related physical strain in predisposed individuals increases their risk of progression to RA, an effect mediated by more severe subclinical joint inflammation (Kronzer et al., 2022; van Dijk et al., 2024; Zeng et al., 2017) whereas alleviation of joint mechanical loading prevents the development of experimental arthritis in mice (Cambré et al., 2018). In addition, forces such as synovial fluid viscosity (Kofoed and Barceló, 1978; Tamer, 2013) and changes in synovial tissue stiffness due to extracellular matrix remodelling (Wei et al., 2023) can also impose mechanical stress on the cells. However, how the synovial tissue cells sense and respond to these various mechanical stimuli has not been explored.

PIEZO1 is a specialized mechanosensitive ion channel that detects changes in the biomechanical environment of cells (Coste et al., 2010). Its activation occurs when the cell membrane is stretched in response to a wide variety of extracellular and intracellular forces (Botello-Smith et al., 2019; Coste et al., 2010). It has been shown that PIEZO1 can regulate macrophage biology, such as polarization, phagocytosis, and migration, in response to changes in the tissue environment (Atcha et al., 2021a; Atcha et al., 2024; Atcha et al., 2021b; Geng et al., 2021; Simakou et al., 2021; Solis et al., 2019; Tang et al., 2023; Wang et al., 2024). However, whether PIEZO1 is expressed by synovial tissue resident macrophages and whether its activation contributes to changes in STM phenotype and the onset of inflammation in the joint remain unknown. In this study, we used synovial biopsies, single cell transcriptomics and animal models to establish the role of PIEZO1 in regulating joint immune biology.

## Results and Discussion

### PIEZO1-activated macrophages acquire a phenotype resembling the inflammation permissive TREM2^low^ lining layer STMs

First, we sought to map PIEZO1 expression in healthy human synovial tissue. Using a combination of anti-CD68 (macrophage marker) antibody staining and PIEZO1 RNAscope, we observed that PIEZO1 was abundantly expressed in lining-layer (LL) cells, namely LL STMs that form a barrier-like structure, and in LL synovial fibroblasts beneath them, whereas PIEZO1 expression was scarce in the sublining layer (SL) **(Fig. 1A).** This indicates that PIEZO1 may play a specific role in regulating lining-layer biology, including that of STMs. In our recent work, we showed that all human synovial tissue-resident macrophages, including lining-layer TREM2^pos^VSIG4^pos^MERTK^pos^ STMs, can originate from monocyte precursors, unlike in mice, where they arise from prenatal precursors (Elmesmari et al., 2025). Therefore, to mimic the *in vivo* system, we used monocyte-derived macrophages (MDM) and stimulated them with the PIEZO1-specific chemical agonist Yoda1 at an optimised dose of 15–20 µM (**Fig. S1A-D).** This agonist binds to mechanosensitive regions of the channel and induces its opening in a manner analogous to mechanical force, thereby mimicking *in vivo* conditions (Botello-Smith et al., 2019; Kinsella et al., 2024; Syeda et al., 2015). Monocyte-derived macrophages expressed PIEZO1 **(Fig. 1B)**, and their activation with Yoda1 resulted in an immediate increase in Ca²⁺ signalling **(Fig. 1C)** as compared with the DMSO control, indicating PIEZO1 channel activation, thus validating our system. To establish the role of PIEZO1 in defining lining-layer STM phenotypes, we stimulated monocyte-derived macrophages (n = 8 healthy donors) with Yoda1 for 12 h to capture the immediate response to PIEZO1 activation and for 72 h to capture the phenotypes that emerge upon continuous PIEZO1 activation, using single-cell transcriptomics **(Fig. 1D and Fig. S1E-G).** As expected, control (DMSO-treated) macrophages formed a transcriptionally uniform population (FOLR2^pos^ cluster, **Fig. 1E**), characterised by high expression of the pan-macrophage marker *FOLR2* and tissue-resident macrophage markers associated with immune-regulatory functions, namely *TREM2* (Jaitin et al., 2019) and *VSIG4* (Li et al., 2017), whose activation limits the inflammatory response of macrophages. In addition, we observed an MMP9^pos^ cluster representing an M-CSF–driven remodelling phenotype (Lepidi et al., 2001), a small fraction of undifferentiated monocytes (FCN1^pos^ cluster) and a population of proliferating macrophages (clusters 5, 7, and 10). Activation of PIEZO1 significantly transformed this pan M-CSF macrophage phenotypes over time **(Fig. 1D-H and Fig. S1E-F).** PIEZO1 activation for 12 h led to rapid changes in macrophage phenotype, including rewiring of the transcriptional regulon, including increased expression of multiple transcription factors, including *HIF1α, NR1H3* (encoding LXRα), *PPARG,* and *NR3C1* (encoding the glucocorticoid receptor). These changes were associated with the emergence of a cluster characterised by decreased expression of the immune-regulatory receptors *TREM2* and *VSIG4,* increased expression of pro-inflammatory cytokines (e.g. *IL-1β*) and chemokines (e.g. *CXCL8* and *CCL3*), and upregulation of phagocytic receptors such as *MERTK* and *CD36*, as well as integrins and proteins essential for phagosome function (the VSIG4^low^MERTK^high^CD36^high^cluster). Continuous 72h PIEZO1 activation resulted in progression of this phenotype into multiple states, including a cluster with high expression of MHC class II molecules (HLA^high^ cluster), a cluster enriched in lipid-metabolic pathways (APOE^pos^APOC1^pos^ cluster), and a phenotype expressing high levels of genes associated with extracellular matrix and bone-remodelling pathways (SPP1^pos^ cluster) **(Fig. 1D–H and Table S1-3).** To gain deeper insight into the signature induced by PIEZO1 activation, we also performed bulk whole transcriptome RNAseq on Yoda1-activated macrophages (n = 6) at 12 h **(Table S4, Fig. S1H-I).** Pathway analysis on these data indicated that PIEZO1 activation limits the proliferation of myeloid cell precursors and drives their tissue terminal differentiation, for example promoting cell–to-cell adhesion pathways **(Fig. S1H),** including tight junction proteins **(Fig. S2A),** which are unique features of lining-layer STMs (Culemann et al., 2019). Validation of these findings at the protein level was performed in macrophages derived from inducible pluripotent stem cells (iPSCs), which more faithfully recapitulate tissue remodelling programs as compared with MDMs (Rajab et al., 2021). PIEZO1 activation in these macrophages significantly increased the expression of the lining-layer tight junction protein TJP1 **(Fig. S2B).** Thus, to determine whether the emerging PIEZO1-triggered macrophage phenotypes are related to lining-layer STM subsets, we generated a PIEZO1 transcriptomic signature comprising 301 upregulated and 199 downregulated genes, identified across our scRNAseq and bulk RNAseq datasets of macrophages treated with a PIEZO1 agonist for 12h **(Table S5).** This signature was used to score synovial-tissue myeloid cell clusters in a large single-cell synovial tissue dataset (MacDonald et al., 2024b) **(Fig. 1I–J and Table S5).** Visualisation of the PIEZO1 transcriptomic signature score (0 = lowest, 10 = highest) on the synovial-tissue myeloid-cell UMAP **(Fig. 1K)** revealed that PIEZO1-activated macrophages closely map to the phenotypes of lining-layer STMs, namely TREM2^low^SPP1^pos^ and SPP1^pos^ clusters, which characterise the hyperplastic lining-layer of inflamed joint of active RA (Alivernini et al., 2020b; Elmesmari et al., 2025; Hanlon et al., 2024; Zhang et al., 2023) and Juvenile Idiopathic Arthritis (JIA) (Bolton et al., 2025), but not to the joint-protective TREM2^pos^VSIG4^pos^ homeostatic phenotype found in healthy joints. In addition, consistent with the cytokine signature induced by PIEZO1 activation, the PIEZO1-specific signature also mapped to the chemokine-high, TNF-high STM cluster (ICAM1^pos^), which is present in active RA (Alivernini et al., 2020b; Elmesmari et al., 2025). Violin-plot visualisation of the PIEZO1-upregulated gene score across myeloid-cell clusters confirmed this pattern **(Fig. 1L).**

**Figure 1.**
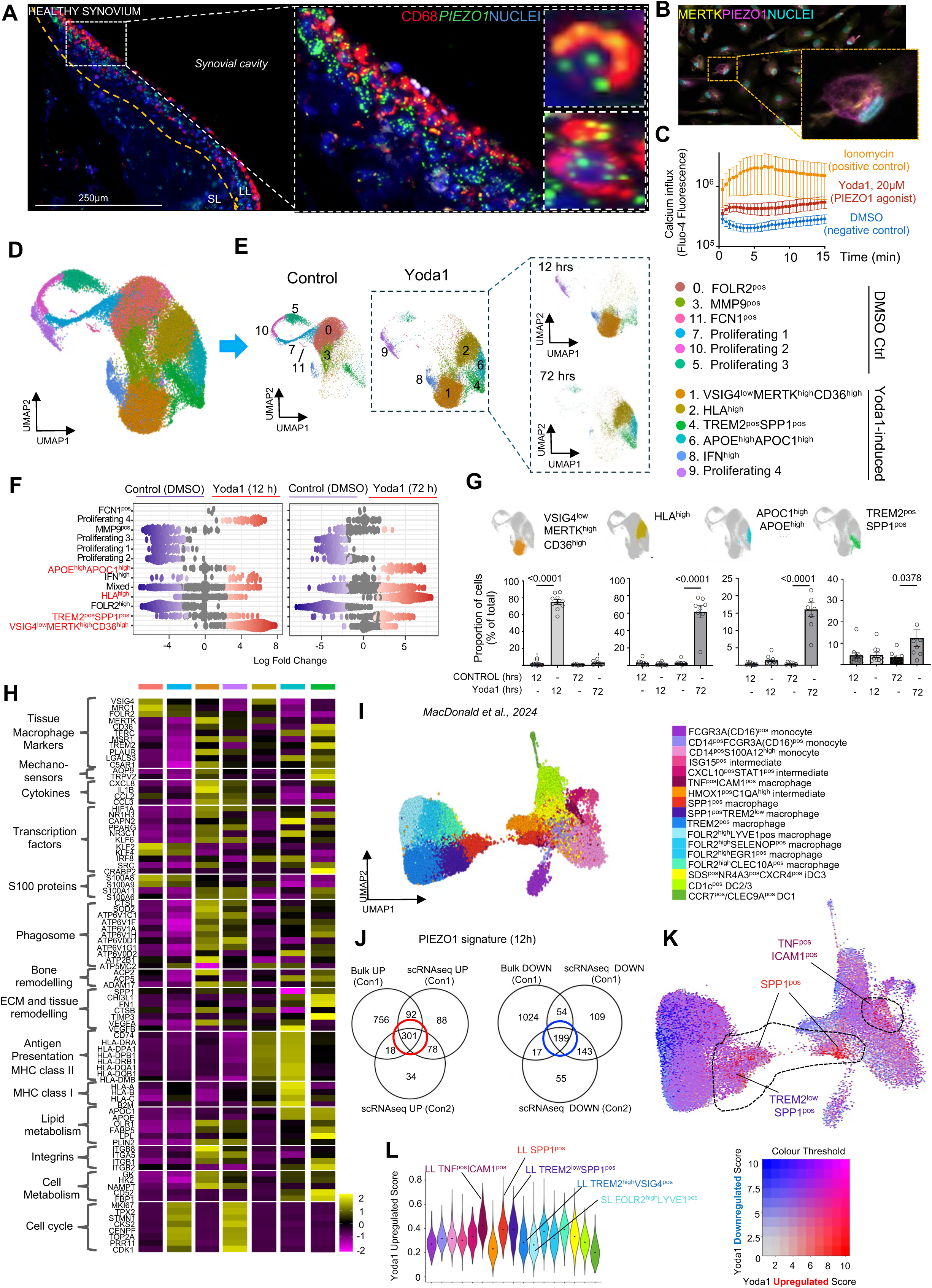
Legend. PIEZO1 activated macrophages acquire phenotype resembling inflammation permissive TREM2^low^ lining-layer STM phenotype. **(A)** mRNA expression of *PIEZO1* (green) in CD68^pos^ (red) macrophages in the healthy synovium. Nuclei stained with SYTO-83 (blue). Representative of n=2 healthy donors. LL = synovial lining-layer; SL = sublining layer; orange dotted line denotes the border of the lining-layer. **(B)** Protein expression of PIEZO1 (magenta) in monocyte-derived macrophages (MDM) cultured in media with M-CSF (20 ng/ml) for 6 days, expressing the tissue-resident marker MERTK (yellow). Nuclei stained with DAPI (cyan). Representative of n=2 healthy donors. Additional data in Figure S2C. **(C)** Change in fluorescence (Fluo-4 dye) to demonstrate calcium influx in monocyte-derived macrophages (MDM) following Yoda1 stimulation (20 µM, red) across time = 0 to 10 minutes. Positive control Ionomycin (orange) and negative control DMSO (blue) included. **(D)** UMAP visualisation of single cell transcriptomics of monocyte-derived macrophages (MDM, n=8 donors) treated with DMSO (control) and Yoda1 (20μM) for 12 and 72 hours (40805 cells, clustering dimensions 25, resolution 0.5) conditions across 2 experiments. Additional information in Table S1 and in Methods. **(E)** Split UMAP of annotated clusters as in (D) between conditions: Yoda1 or DMSO stimulation at overnight (12-hour) and 72-hour timepoints, respectively. **(F)** Differential abundance showing changes in different clusters frequencies between conditions. Data are visualized as MiloR neighbourhood graphs, where nodes represent neighbourhoods, coloured by their log fold change across conditions. Neighbourhoods with non-differential abundance (FDR > 10%) are coloured grey, and node size reflects the number of cells in each neighbourhood. Colours represent upregulated (red) and downregulated (violet) neighbourhoods in Yoda1 samples compared to DMSO at 12 hours (left) and 72 hours (right). **(G)** Proportion plots of Yoda1-induced clusters from (D), for overnight (12 hours) and 72 hours, respectively. Data presented as bar-plot with SD of the mean. Each dot represents a donor (N=8), Statistical analysis performed using one-way Anova with Tukey’s correction for multiple comparison. Exact p values are on the graph. **(H)** Pseudobulk heatmap of markers of the selected clusters identified in (D) (FindAllmarkers, logfc.threshold 0.25). Only the clusters of interest from Yoda1-stimulated conditions, and the control FOLR2^pos^ and proliferating 1 clusters shown. Genes grouped for functionality. **(I)** UMAP visualisation of published dataset from *MacDonald et al., 2024 (DOI: 10.1016/j.immuni.2024.11.004)*, showing myeloid subsets identified in synovial tissue biopsies from healthy donors (n=6), patients with active RA(n=18), and in RA disease remission (n=7) **(J)** Integration of the top upregulated (301) and top downregulated (199) genes in MDM cultured with 20 ng/ml for 5 days and then stimulated with Yoda1 for 12 hours across two scRNA-seq conditions (Con1=Condition1, where media supplemented with fresh M-CSF was replenished before adding Yoda1, n=6 and Con2-Condition2 where media was not replenished before adding Yoda1, n=2) and bulk RNA-sequencing dataset (Con1, media not replenished, n=6). Data are presented as a Venn diagram showing PIEZO1-induced transcriptional changes identified by bulk RNA sequencing (adjusted p < 0.05 and log2 fold change > 0.58) and single-cell RNA sequencing (adjusted p < 0.05 and log2 fold change > 0.25) in overnight-stimulated cells. **(K)** Visualisation of the PIEZO1 transcriptomic signature score (0 = lowest, 10 = highest) on the synovial-tissue myeloid-cell UMAP as in (I) using STACAS. Genes from (J) were used to generate PIEZO1 signature. **(L)** Violin plot illustration of PIEZO1 transcriptomic signature on the scale from 0=the lowest to 1= the highest in different synovial tissue myeloid cell clusters from (I).

We next validated the PIEZO1-induced inflammation-permissive TREM2^low^ phenotype at the protein level. In agreement with the transcriptomic signature **(Fig. 2A),** we observed a time-dependent decrease in the percentage of VSIG4⁺ macrophages and in the expression levels (MFI) of both VSIG4 and TREM2 upon PIEZO1 activation. This was accompanied by increased expression of antigen-presenting molecules (MHC II) and the phagocytic receptors MERTK and CD36, suggesting enhanced tissue antigen clearance and antigen presentation **(Fig. 2B–D and Fig. S2C).** We observed that both monocyte- and iPSC-derived macrophages showed increased sensitivity to PIEZO1 regulation **(Fig. 2B)**, extending to a significant downregulation of the additional homeostatic lining-layer STM marker CD206 when cells were matured with a low dose of M-CSF, which more closely reflects local tissue concentrations **(Fig. S2D–F).** Investigation into the functional soluble mediators regulated by PIEZO1 activation revealed a significant increase in the mRNA expression of multiple chemokines that regulate neutrophil and monocyte migration. Consistent with predictions from both bulk transcriptomic **(Fig. 2E)** and scRNAseq data (Fig. 1H), PIEZO1 stimulation triggered the release of the neutrophil-attracting chemokine IL-8 to levels comparable to those induced by TLR engagement **(Fig. 2F)**. This finding suggests a unique role for PIEZO1 in neutrophil recruitment in response to aberrant mechanical stimuli that is independent from other triggers such as TLRs, thereby potentially initiating the early steps of inflammation in the joint. Furthermore, PIEZO1 activation significantly enhanced TLR-primed production of the monocyte-attracting chemokine CCL3 **(Fig. 2G).** Recent data from experimental arthritis models has established that chemokines released specifically by the lining-layer STMs represent a critical first step in localizing inflammation within the joints (Zec et al., 2023b). Thus, our data suggest that lining-layer STMs in joints can sense biomechanical stimuli via PIEZO1, and that PIEZO1 activation may induce their inflammation-permissive phenotype by triggering chemokine release, reducing the expression of regulatory receptors, and increasing antigen-presenting capacity, thereby initiating joint inflammation.

**Figure 2.**
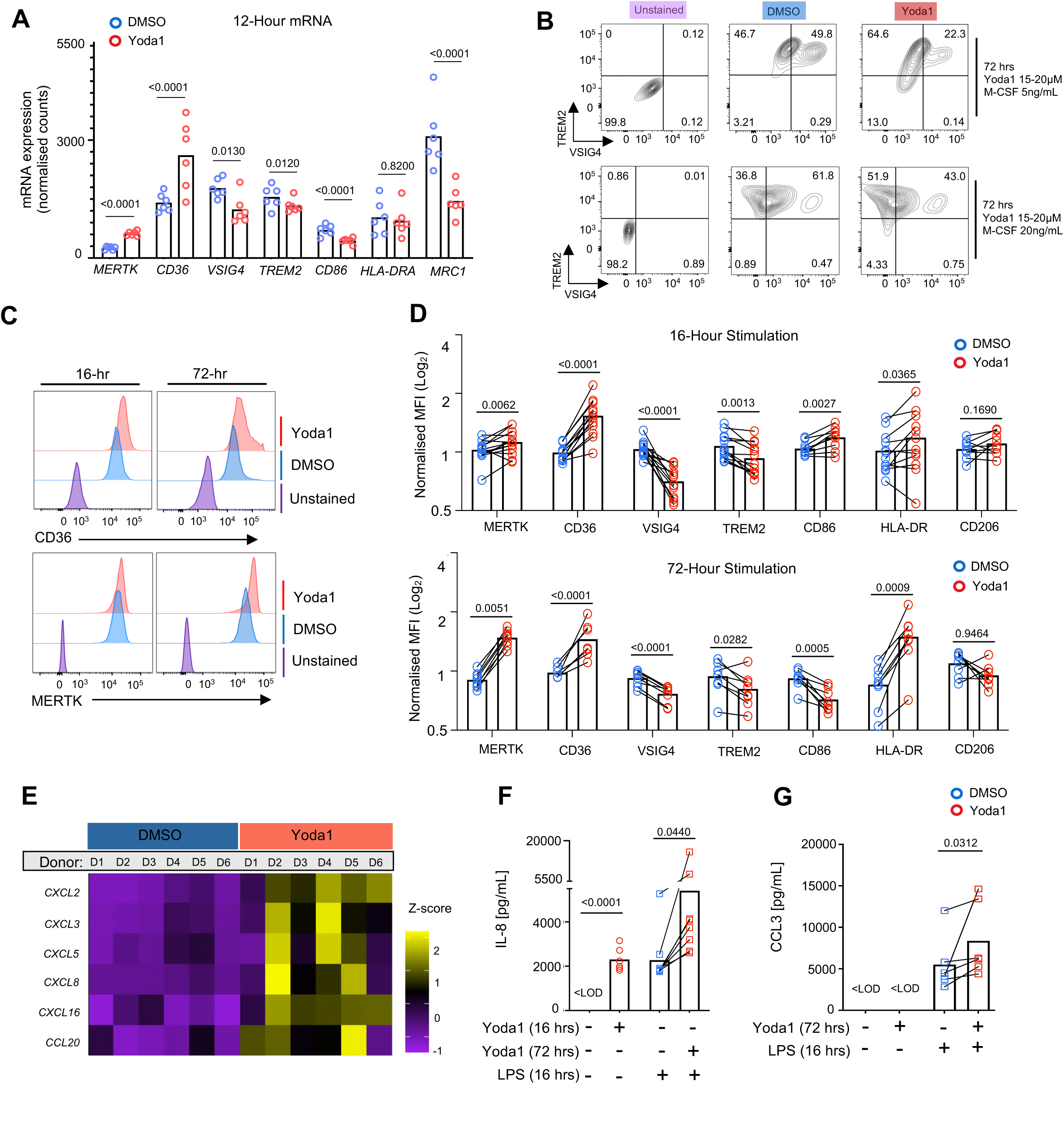
Legend. PIEZO1 activation triggers chemokine productions by macrophages. **(A)** mRNA expression (bulk RNAseq) of selected lining-layer STMs markers in monocyte-derived macrophages stimulated with Yoda1 (20 µM) or control DMSO (0.1%) for 12 hours. Each dot represents one donor, N=6 healthy donors. DEseq2 with adjustments for multiple comparison. Exact p values are on the graph. **(B)** Representative flow cytometry of TREM2 and VSIG4 expression by macrophages derived from monocytes with reduced (5 ng/ml) or standard (20 ng/m)l M-CSF and treated with Yoda1 (15-20µM) for 72h. **(C-D)** Median fluorescence intensity (MFI) of selected lining-layer STM markers in monocyte-derived macrophages stimulated with Yoda1 or control DMSO (0.1%) for 16 and 72 hours. (C) Representative histograms of CD36 and MERTK. (D) MFI of the STM markers normalised to MFI of macrophages in media with M-CSF alone. Each dot represent one donor, N=8 healthy donors across at least 3 different experiments. Paired T test, exact p values provided on the graphs. **(E)** Heatmap illustrating mRNA levels of chemokines upregulates in macrophages upon overnight stimulation with 20 µM Yoda1. Data normalised for inter-quartile range. Colour intensity indicates row scaled (z-score) gene expression with purple as low and yellow as high. Bulk RNA sequencing; DEseq2 with correction for multiple comparison genes with adjusted p-value (<0.0001) are shown. **(F-G)** IL-8 (F) and CCL3 (G) release by monocyte-derived macrophages stimulated upon PIEZO1 activation. Cells were incubated with with control DMSO (0.1%) (blue) or 20 µM Yoda1 (red) in the absence or presence of LPS (1ng/mL) at different time points. LOD = limit of detection. Each dot represents one donor, N=6) across 5 experiments. Paired T test, exact p values provided on the graphs.

### PIEZO1 activation in the mouse joint triggers inflammation

To test the hypothesis that PIEZO1 activation in lining-layer STMs contributes to the localisation of inflammation in joints, and thereby to arthritis development and severity, we first evaluated PIEZO1 expression in STMs across the RA disease trajectory (Alivernini et al., 2020a) **(Fig. 3A)**. Single-cell transcriptomic analysis confirmed that PIEZO1 is predominantly expressed in the lining-layer clusters. The highest expression levels, as well as the largest proportion of PIEZO1-expressing cells within a cluster, were observed in the pathogenic lining-layer phenotypes (e.g., the SPP1^pos^ cluster), consistent with their high PIEZO1-regulated signature score, as deconvoluted in Fig. 1J–K. This phenotype of lining-layer STMs dominates the hyperplastic lining-layer of patients with active RA, with abundance significantly reduced when sustained disease remission is achieved **(Fig. 3A**) (Alivernini et al., 2020a; Elmesmari et al., 2025). Thus, this suggests that the lining-layer becomes increasingly receptive to changes in the biomechanical environment as it becomes inflamed and hyperplastic **(Fig.3A).** Spatial deconvolution using RNAscope and antibodies against PIEZO1 **(Table S6)** combined with macrophage staining, confirmed an increased PIEZO1 mRNA/protein copy number per lining-layer macrophage in active RA compared with lining-layer cells from patients in sustained remission with healthy, tissue-like synovial structure **(Table S7).** It also showed that the PIEZO1 mRNA copy number per cell, in lining-layer STMs from individuals at-risk of RA, is similar to that observed in active RA, suggesting that these cells may already be poised for an enhanced response to mechanical stimuli **(Fig. 3B–E and Fig. S2F-H).** Altogether, this confirms a potential role for biomechanical stimuli-mediated STM activation in driving joint immune responses.

**Figure 3.**
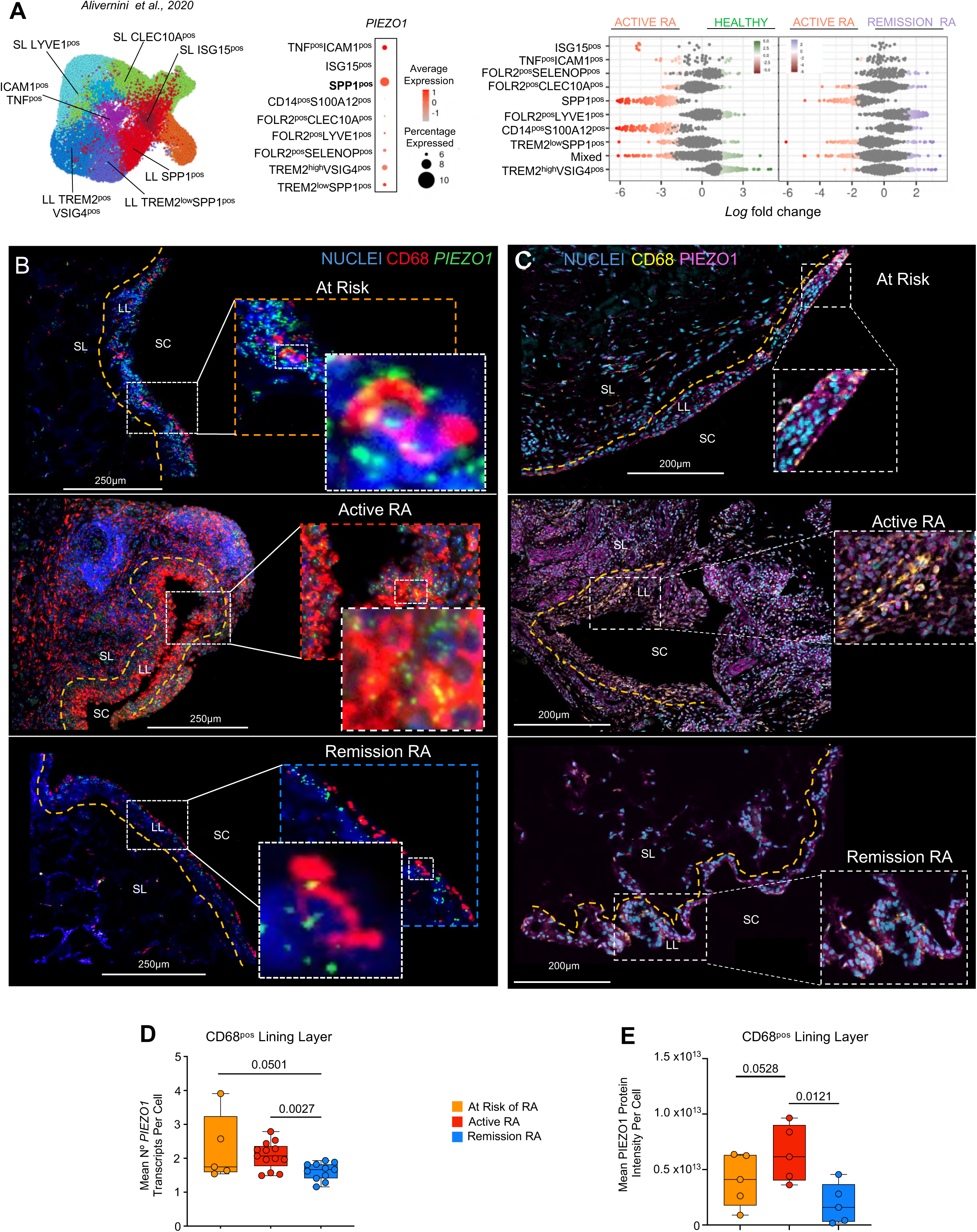
Legend. PIEZO1 is highly expressed by macrophages in hyperplastic lining-layer of active RA. **(A)** Expression of *PIEZO1* mRNA in STM clusters across different joint conditions (Alivernini *et al*., 2020, *DOI: 10.1038/s41591-020-0939-8*), showing the highest expression of *PIEZO1* in the lining-layer SPP1^pos^ STM clusters characteristic of active RA. Data is presented as (i) a dot plot showing average and percentage log-normalised expression of PIEZO1 in synovial macrophage clusters and (ii) differential abundance in major macrophage clusters frequencies between health (n=4), active RA (n=12) and RA in remission (n=6). Data are visualized as MiloR neighbourhood graphs, where nodes represent neighbourhoods, coloured by their log fold change across conditions. Neighbourhoods with non-differential abundance (FDR > 10%) are coloured grey, and node size reflects the number of cells in each neighbourhood. Colours represent neighbourhood upregulated in healthy (green), active RA (red) and remission RA (violet) **(B)** Expression of *PIEZO1* mRNA (green) evaluated by RNA Scope in lining layer CD68-positive (red) macrophages in synovial tissue biopsies of RA patients at different stages of disease: at risk (top); naïve to treatment (centre); remission (bottom). Inset images depict higher magnification of individual cells, showing colocalization of *PIEZO1* transcripts and CD68^pos^ macrophages in the lining-layer. Nuclei stained with SYTO-83 (blue). LL = lining-layer; SL = sublining; SC = synovial cavity; orange dashed line indicates the border of the lining layer. Scale bar = 250µm. **(C)** Expression of PIEZO1 protein (magenta) in CD68-positive (yellow) macrophages in synovial tissue biopsies of RA patients at different stages of disease: at risk (top); naïve to treatment (centre); remission (bottom). Inset images depict higher magnification of lining-layer cells. Nuclei stained with DAPI (cyan). LL = lining-layer; SL = sublining; SC = synovial cavity; orange dashed line indicates the border of the lining layer. Scale bar = 250µm. **(D)** Quantification of *PIEZO1* mRNA transcripts per cell in LL CD68^pos^ macrophages in patients at risk of RA (N=2), with active RA (N=4) or remission RA (N=4). Each dot represents one FOV from synovial biopsies, and the expression is determined by number of transcripts per FOV. Unpaired T test, p values are provided on the graph, ST = synovial tissue. **(E)** Quantification of PIEZO1 protein in LL CD68^pos^ macrophages. Quantification was performed for each CD68 positive cells, and raw IntegratedDensity of PIEZO1 measured in the single channel of its fluorophore using ImageJ. Data represents the mean of corrected total cell fluorescence (CTCF) of multiple cells per each tissue (N=5 per condition); Unpaired T test, p values are provided on the graph.

Thus, to test if activation of PIEZO1 in the joint can trigger inflammation, we used a model of intra-articular Yoda1 injection to mimic biomechanical stimulation. We confirmed that, as in human synovium, Piezo1 is expressed in the healthy mouse lining-layer **(Fig.4A).** To minimise variability in joint biomechanical activity between mice, we injected Yoda1 at a dose optimised for the *in vivo* model [5 μM, (Pinto et al., 2025)] into the right knee and vehicle control (DMSO) into the left knee of individual healthy mouse at days 0 and 3 to mimic prolonged Piezo1 activation (n=8 mice altogether). We showed previously that the uptake of mediators delivered via intra-articular injection is mostly limited to synovial lining-layer and synovial cartilage cells (Huesa et al., 2016). To capture both the immediate and follow up consequences of prolonged Piezo1 activation, we assessed changes in the synovium at day 4 and day 7 **(Fig. 4B).** Histological analysis **(Fig. 4C**) and flow cytometry of digested synovial tissue **(Fig. 4D-F and Fig. S3B)** from contralateral joints showed that activation of Piezo1 (two injections of Yoda1, two days apart) triggers lining-layer hyperplasia and changes in the phenotype of lining-layer macrophages (Cx3cr1^pos^ STMs), similar to those observed in the human *in vitro* model, including increased expression of Mhc II and Mertk. In addition, Piezo1 activation in joints led to a significant influx of early inflammatory cells, neutrophils and monocytes, into the synovium at day 4 as compared with control contralateral joints (**Fig. 4G-I).** These findings are consistent with the release of neutrophil and monocyte chemoattractants and activators, IL-8 and CCL3 from macrophages upon PIEZO1 activation, as shown in Figure 2. This inflammation was largely transient; by 96 hours after last Yoda1 injection (experimental day 7), almost no immune infiltrates remained in the synovium. This may be attributable to a Piezo1-triggered negative feedback mechanism on inflammation, reflected by high expression of efferocytosis receptors (Fig. 2C-D and Fig.4F), including Mertk, which possess strong inflammation-limiting properties in STMs (Alivernini et al., 2020b), thereby preventing the development of chronic inflammation once the aberrant mechanical stimulation ceases. This suggests that development of inflammation-permissive, chemokine-producing lining-layer STMs may require prolonged perturbation of the mechanical environment, for example at the cellular level through age-associated alterations in extracellular matrix stiffness, or at the organ level through increased and repetitive mechanical strain.

**Figure 4.**
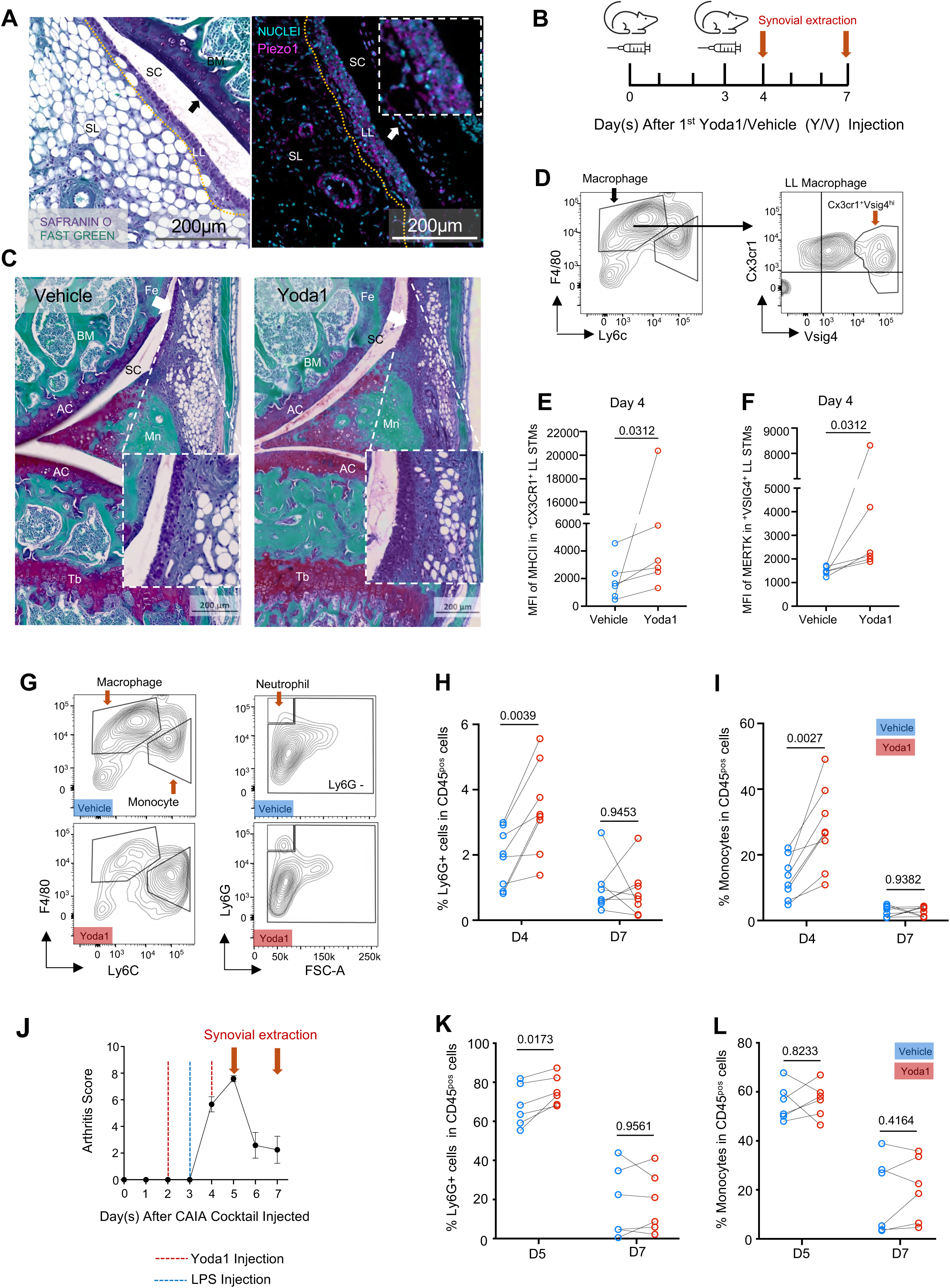
Legend. Piezo1 activation triggers joint inflammation. **(A)** Histological staining of collagen (purple), bone and ligaments (green), and soft tissues and nuclei (H&E) in the healthy mouse knee joint (left), and expression of Piezo1 (magenta) with nuclei (cyan) in healthy mouse synovium (right). Piezo1 was expressed in the synovial lining-layer (LL), and to a lesser degree in sublining (SL) endothelium and articular cartilage chondrocytes. SC = synovial cavity; BM = bone marrow. The arrow indicates chondrocytes in the articular cartilage opposite the synovial LL and separated by the SC; the orange dashed line indicates the border of the LL. **(B)** Schematic diagram of experimental timeline showing contralateral intra-articular injection of Piezo1 agonist Yoda1 (5µM, right knee) or vehicle (0.025% DMSO, left knee). **(C)** Histological representation of mouse synovium (purple), bone and ligaments (green), and soft tissues and nuclei (H&E) following injection with vehicle (left) and Yoda1 (right) injected knees, stained with haematoxylin & eosin, safranin-O/fast green. White arrow = lining-layer. Collagen (purple); bone and ligament (green); soft tissues and nuclei (H&E). Images representative of at least N=3 mice. **(D)** Representative flow cytometry gating strategy for mouse lining layer Cx3cr1+Vsig4+ macrophages Cells that were Ly6g^neg^Cd3^neg^Cd45^pos^ were gated for F4/80 and Ly6c expression. F4/80^pos^ that were negative for Ly6c and positive for Cx3cr1 and Vsig4 were identified as lining-layer STMs. **(E-F)** MFI of Mertk **(E)** and I-A/E **(F)** in lining-layer Cx3cr1+1Vsig4hi STMs, assessed by flow cytometry on Day 4, 24 hours post second intra-articular injection. Each dot represents one mouse, paired values from vehicle and Yoda1 injected knees, n = 6 mice across two experiments; statistical analysis performed using Wilcoxon test. **(G)** Representative flow cytometry of monocytes and macrophages (left) as well as neutrophils (right) in synovial tissue of knees injected with vehicle (top) or Yoda1 (bottom) from the same mouse, 24 hours post second intra-articular injection. **(H-I)** Percentage of neutrophils **(H)** and monocytes **(I)** in total Cd45^pos^ synovial tissue cells in vehicle (blue)- and Yoda1 (red)-injected knees on Day 4 and Day 7. Paired values from vehicle and Yoda1 injected knees, n = 8 mice across two experiments; statistical analysis performed using Wilcoxon test. **(J)** Arthritis score of mice with collagen antibody induced arthritis (CAIA) and schematic showing Yoda1 injection timeline. Each mouse had Yoda1 injected in its right knee and vehicle in its left knee on Day 2 and Day 4 after IV injection of CAIA antibody cocktail. LPS was injected IP on Day 3. Synovium was extracted on Day 5 and Day 7. **(K-L)** Percentage of neutrophils **(K)** and monocytes **(L)** in total Cd45^pos^ synovial tissue cells in vehicle (blue)- and Yoda1 (red)-injected knees on Day 5 and Day 7 as described in **(J).** Paired values from vehicle and Yoda1 injected knees, n = 8 mice across two experiments; statistical analysis performed using Wilcoxon test.

Having established increased PIEZO1 expression in lining-layer STMs in active RA, we next sought to investigate whether PIEZO1 contributes to the severity of established arthritis. We utilised the collagen antibody-induced arthritis (CAIA) model and performed intra-articular injections of Yoda1 or vehicle at two time points: before the onset of clinical symptoms (day 2) and before the peak of arthritis (day 4) **(Fig. 4J).** Similar to the phenotype observed in arthritis-naïve mice described above, we found that Piezo1 activation in the joints increased neutrophil recruitment triggered by collagen immune complexes at the peak of arthritis (day 5) **(Fig. 4K),** compared with vehicle-treated controls. There was no difference in monocytic infiltration between the Yoda1- and vehicle-injected groups **(Fig. 4L),** suggesting that mechanical signal may be a key driver of neutrophil influx into inflamed joints while redundant for monocyte recruitment. In summary, aberrant activation of PIEZO1 in joints, whether driven by continuous biomechanical stimuli and/or increased expression of this ion channel may be responsible for promoting the localisation of inflammation, thereby contributing to disease onset and subsequently to arthritis severity.

### PIEZO1 driven inflammation permissive macrophage phenotype is mediated by calpains and HIF-1α

Next, we sought to define the downstream signalling pathways of PIEZO1 that drive the development of the TREM2^low^VSIG4^low^MERTK^high^CD36^high^ macrophage phenotype, which resembles the inflammation-permissive lining-layer STM clusters. We showed that PIEZO1 activation with Yoda1 results in immediate Ca2+ signalling which spikes for 4 minutes (Fig.1C). PIEZO1-triggered Ca2+ entry to the cells mediates actin cytoskeletal remodelling via activation of calpain, a calcium-dependent cysteine protease (Baratchi et al., 2024; Liu et al., 2018a; Liu et al., 2018b; Potter et al., 1998). We demonstrated that upon engagement of PIEZO1, F-actin polymerization changed rapidly in monocyte-derived macrophages (1 minute after Yoda1 stimulation) **(Fig. 5A-B).** This resulted in alterations in nuclear size and shape, indicating active cytoskeletal remodelling over time **(Fig. 5C).** We also observed a PIEZO1-driven increase in spindle-shaped macrophages **(Fig. 5D and Fig. S3C),** a characteristic feature of cellular responses to changes in their immediate mechanical environment (Jetta et al., 2021; Rashidi et al., 2025) further validating our *in vitro* system for dissecting PIEZO1 signalling. Recently, in addition to activating calpain, PIEZO1 has been shown to stabilize HIF1α (Solis et al., 2019), a transcription factor that integrates macrophage responses to diverse environmental stimuli (McGettrick and O’Neill, 2020). Therefore, we tested the contribution of both calpain and HIF1α pathways to the development of an inflammation-permissive macrophage phenotype. We observed that PIEZO1 increased *CAPN2* mRNA at 12 hours after Yoda1 stimulation **(Fig. 5E).** Incubation of macrophages with the calpain inhibitor PD150606 (Low et al., 2014) during the 12-hour period of PIEZO1 activation revealed that calpain inhibition prevented at least some aspects of the PIEZO1-induced macrophage phenotype, such as the upregulation of CD36 **(Fig. 5F-5G).** We next looked at the role of HIF1α. We established that HIF1α was upregulated in macrophages stimulated with Yoda1 for 12 hours, at both the mRNA **(Fig. 5H)** and protein levels **(Fig. 5I).** Furthermore, cell trajectory and pseudotime analysis of single-cell data from Yoda1-stimulated macrophages showed that *HIF1A* was consistently upregulated throughout the transition from control macrophages to the Yoda1-induced terminally differentiated VSIG4^low^TREM2^low^SPP1^pos^ cluster over time **(Fig. 5J and Fig. S1G)**, supporting its role in the PIEZO1-driven STM phenotype. To test this, we generated HIF1α-deficient monocytes using a CRISPR-Cas9 approach (n = 3). The monocytes were then differentiated into macrophages, followed by stimulation with Yoda1 for 15 hours. *HIF1A* deletion prevented both PIEZO1-driven inhibition of VSIG4 and upregulation of CD36, suggesting a potential role of PIEZO1-induced HIF1α in coordinating the development of the inflammation permissive macrophage phenotype **(Fig. 5K and Fig. S3D).** Using macrophage staining together with digital spatial transcriptomics (GeoMx), we compared *HIF1A* expression in hyperplastic lining-layer macrophages from synovial biopsies of patients with active RA, which is dominated by the inflammation-permissive TREM2^low^SPP1^pos^ and SPP1^pos^ clusters, with lining-layer macrophages from patients who achieved sustained disease remission, which exhibit reinstated homeostatic STMs (Elmesmari et al., 2025). Consistent with the PIEZO1 signature of hyperplastic lining-layer STMs clusters in active RA (TREM2^low^SPP1^pos^ and SPP1^pos^), these macrophages showed significantly higher expression of *HIF1A* compared to lining-layer macrophages in disease remission **(Fig. 5L and Table S8).** In summary, HIF1α and calpains, at least in part, may contribute to the development of the PIEZO1-induced inflammation-permissive STM phenotype underlying RA pathology.

**Figure 5.**
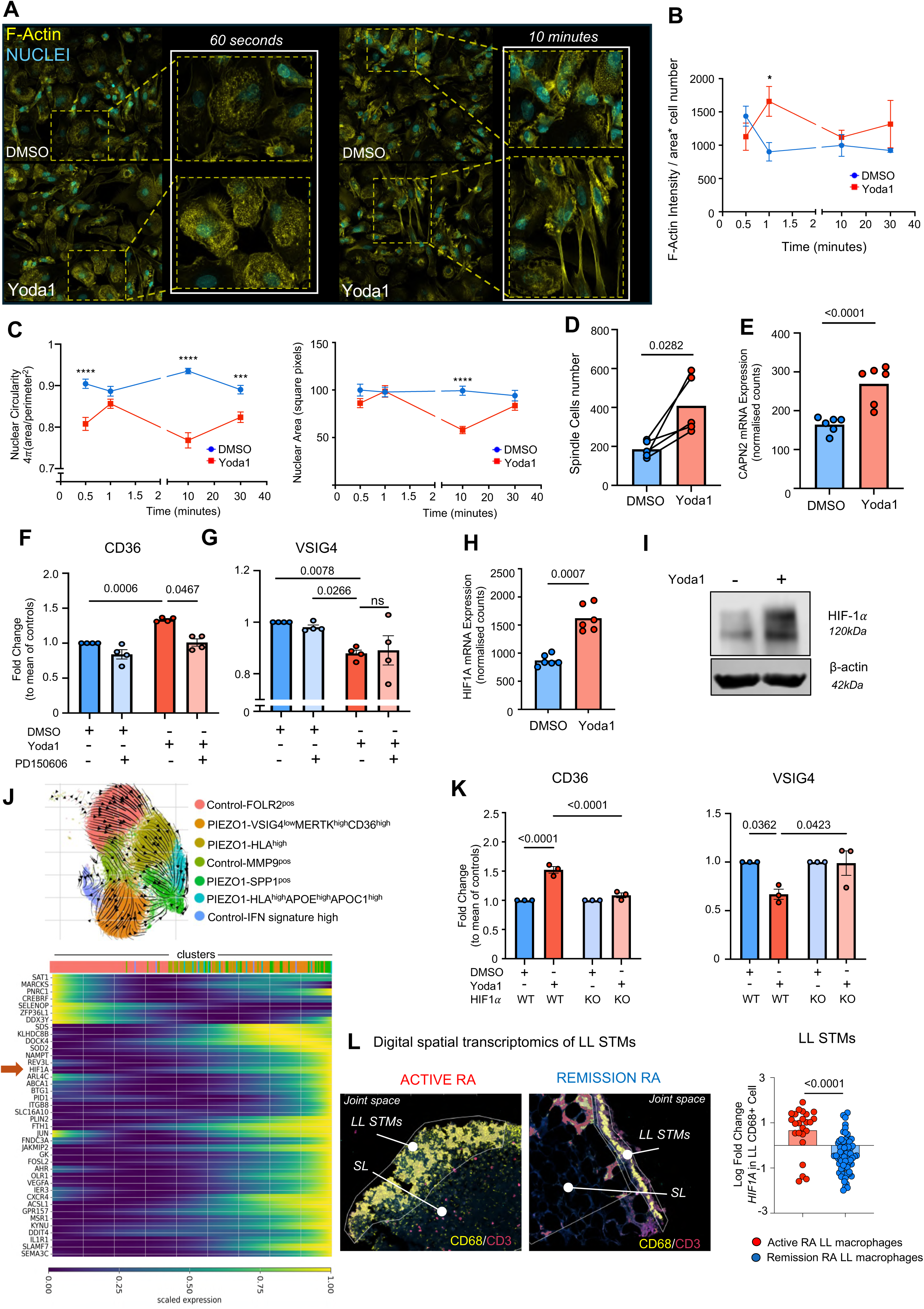
Legend. PIEZO1 activation drives inflammation permissive macrophage phenotype through Calpain and HIF1α signalling. **(A)** F-actin visualisation in phalloidin-stained macrophages stimulated with 20µM Yoda1 or control 0.1% DMSO for 30 seconds, 1 minute, 10 minutes and 30 minutes. Images are representative of N=3 experimental repeats. Images taken at 63X magnification as Z stack performed at 63x magnification and displayed as projections using Zeiss Zen Black software. **(B)** Quantification of F-actin in macrophages per field of view in 3 experimental repeats (N=3) for each timepoint. F-actin was quantified as follows: Intensity= RawIntegratedDensity /(Area/number of cells), using ImageJ software. Number of cells was determined by counting the nuclei in the selected area. Data shown as mean +/- SEM; Unpaired T test, *p≤0.05. **(C)** Comparison of nuclear circularity (left) and size (right), in macrophages stimulated with Yoda or vehicle. The nuclei were identified by positive staining of DAPI. Circularity was measured as 4π(area/perimeter^2), using Image J. Data shown as mean +/- SEM of 45 nuclei selected randomly from technical replicates. Unpaired T test, ****p≤0.0001, ***p≤0.001. **(D)** Number of Spindle macrophages in cultures stimulated with 20µM Yoda1 or 0.1% DMSO for 72 hours. N=5, each dot represents one donor. Grayscale-converted images were pre-processed using Gaussian Blur and thresholding to enhance spindle cell visibility, followed by binary conversion and watershed segmentation to separate adjacent cells. Cell counts were obtained using the Analyze Particles function in ImageJ, applying size and circularity filters (circularity: 0.1–0.5) appropriate for the elongated morphology of spindle cells. Data shown as aligned plot of paired DMSO-Yoda1 cultures from the same donor; statistical analysis was performed using paired T test, exact p value is on the graph. **(E)** Expression of mRNA of *CAPN2* in macrophages stimulated overnight with 20µM Yoda1. Each dot represents one donor (N=6). Data obtained by Bulk RNA sequencing, DEseq2, and p-value (<0.0001) adjusted for multiple comparison. **(F-G)** Expression of CD36 (F) and VSIG4 (G) by Yoda1 stimulated macrophage cultures in the presence of Calpain inhibitor. Macrophages were stimulated with Yoda1 (20µM )or DMSO1 in the presence of Calpain inhibitor PD150606 (100µM) for 12 hours. Data is presented as barplot with the mean of CD36 MFI normalised to the mean of DMSO controls (N=4); statistical analysis performed using repeated-measures one-way Anova, with Tukey’s multiple comparison Test, ns=non statistically significant **(H)** Expression of *HIF1A* mRNA in macrophages stimulated with 20µM Yoda1 overnight as compared to 0.1% DMSO controls. Each dot represents a one donor (N=6). Data obtained by Bulk RNA sequencing, DEseq2, and p-value (<0.0001) adjusted for multiple comparison. **(I)** Representative Westen blot showing HIF-1α protein expression in 20µM Yoda1 or 0.1% DMSO treated cells for 12h. **(J)** Trajectory and pseudotime analysis of single-cell transcriptomics from control and Yoda1-stimulated macrophages. Arrows overlaid on the UMAP, visualising the clusters shown in Fig. 1D, indicate a progression from control macrophage clusters toward a terminally differentiated, PIEZO1-driven SPP1^pos^ macrophage cluster. The pseudotime heatmap highlights genes underlying the transition from a pan-macrophage state to a PIEZO1-driven phenotype, including HIF1A (detailed information in Methods). **(K)** Expression of CD36 and VSIG4 by Yoda1 stimulated wild type (WT) and HIF1α deficient macrophage (KO). WT and KO macrophages were stimulated with Yoda1 (20µM) or DMSO1 for 16 hours. Data is presented as barplot with the mean of CD36 and VSIG4 MFIs normalised to the mean of DMSO controls (N=3); statistical analysis performed using repeated-measures one-way Anova, with Tukey’s multiple comparison Test, exact p values are provided on the graph. **(L)** Digital spatial transcriptomics (GeoMx) analysis of synovial lining-layer macrophages of patients with active RA and in disease remission. *HIF1A* mRNA transcripts were quantified in lining-layer (LL) macrophages across 28 Regions of Interests (ROIs) from synovial biopsies of n=8 RA patients with active disease and 80 ROIs from synovial biopsies of n=16 RA patients in remission. Expression was determined as the number of individual transcripts per ROI. Statistical analysis was performed using Mixed Linear Model (LMM). Each dot represents a single ROI, and the p value is indicated on the graph. Data are presented as the fold change between conditions in normalized *HIF1A* expression relative to the housekeeping gene within each ROI.

In summary, we found that the mechanosensitive ion channel PIEZO1 is highly expressed in the synovial tissue lining-layer, including lining-layer–resident macrophages, indicating that these cells are particularly receptive to mechanical stimuli, including enhanced physical strain that joints can experience. We provide evidence that PIEZO1 activation can lead to a decrease in the regulatory phenotype characteristic of healthy lining-layer STMs and drives, through calpain and HIF1α, a chemokine-releasing, inflammation-permissive phenotype closely related to the TREM2^low^SPP1^pos^ and SPP1^pos^ lining-layer clusters that characterize the hyperplastic lining-layer in RA patients with active disease. Prolonged activation of PIEZO1 in mouse joints triggers joint inflammation, which resolves when the stimuli cease. We speculate that PIEZO1-induced, inflammation-permissive state of lining-layer STMs triggered by aberrant mechanical stress, for example, due to changes in extracellular matrix stiffness or mechanical overload of the joints may be sufficient to localize asymptomatic breaches of immune tolerance in individuals at-risk of RA, thereby triggering the onset of clinical symptoms, and thereafter contribute to disease severity. This is in line with recent findings showing an increased risk of arthritis development with the duration and intensity of work-related strain (van Dijk et al., 2024), and is supported by previous experimental studies in mice showing that, in the absence of biomechanical pressure on the joints, arthritis does not develop (Cambré et al., 2018). Thus, timely interference with PIEZO1 signalling in the joints may be crucial for maintaining and/or reinstating a healthy lining layer in individuals at risk of arthritis, thereby preventing the onset of clinical disease. It is emerging that multiple PIEZO1 allelic variants exist that either increase or decrease PIEZO1 function (Cheng et al., 2025) that confer advantage or disadvantage of different tissues to specific environment (Ma et al., 2021; Nakamichi et al., 2022). Thus, future studies are needed to determine whether specific PIEZO1 variants are responsible for an increased risk of inflammatory arthritis onset under conditions of elevated mechanical load in individuals with a breach of immune tolerance.

## Material and Methods

### Monocyte-derived macrophage cultures

The PBMCs were isolated from blood of healthy donors **(Table S7)** using the density separation with Histopaque-1077 (Sigma-Aldrich, 10771-500ML). The blood was diluted 1:2 with sterile 1X D-PBS (Gibco, 14190-094), and then slowly added on top of Histopaque-1077. Density separation of cells was performed by centrifugation at 2500 RPM for 25 minutes at room temperature, without break. After the centrifugation, the PBMCs were collected from the buffy coat. Total monocytes (StemCell Tech,19058) and CD14 monocytes (StemCell Tech, 19359) were isolated using negative selection as per manufacturer’s protocols. The CD14 positive monocytes were cultured in 24 well plates, at a density of 1.5×10^5 cells per cm^2^ in RPMI-1640 media (Gibco, 21875-034), supplemented with 10% FBS (Sigma-Aldrich, F9665), 2mM L-Glutamine (Sigma-Aldrich, G7513-100ML) and 100 U/mL Penicillin and 100 μ g/mL streptomycin (Gibco, 15140-122), and human M-CSF at 5-20 ng/mL (Peprotech, 30025) for various time points as indicated below.

### The iPSC-derived macrophage (iMacs) cultures

The iPSC cells were kindly provided and then maintained and differentiated into macrophages according to Prof Lesley Forrester’s lab (Lopez-Yrigoyen et al., 2018). Briefly, for iPSC culture, 6 wells plates were coated with 5 µg/mL vitronectin (Thermofisher A31804) at 37 ℃ for 1 hour. The iPSCs were cultured in StemFlex medium (Thermofisher, A3349401) with 10 µM of ROCK inhibitor Y27632 for 24 hours (StemCell Tech, 72302). The following day, media with ROCK inhibitor was removed, and new StemFlex media was added to the cells. The cells were grown until 80% confluency, before passaging. For passaging the iPSC colonies were detached by incubation with the StemMACS Passaging Solution XF (Miltenyi, 130-104-688) for 4 minutes. The cells were then transferred to the new plates and cultured in StemFlex medium. The confluent iPSCs (90%) were then detached in clumps using the EZ passage tool transferred into repellent plates for development of embryonal bodies. The media used for these cultures contained complete StemPRO media (Gibco, A10007-01) supplemented with 2% BSA (Gibco, A10008-01) and BMP4 (50 ng/mL) (Biotechne, 314BP010), VEGF (50 ng/mL) (BioTechne, 293VE010) and SCF (20 ng/mL) (Thermofisher, PHC2115). Cells were fed every other day. After 6-7 days, the embryonal bodies (EB) were visible in suspensions. Embryonal bodies were then cultured in 6-well plates that were coated with 0.1% porcine gelatine (Sigma-Aldrich, G2500). In order to generate the iMacs precursors, embryonal bodies were cultured in serum-free X-Vivo medium (Lonza, 02060Q) with 100 ng/mL M-CSF, 25 ng/mL IL-3 (Peprotech, 200-03), 2 mM L-glutamine (Sigma-Aldrich, G7513), 1% Penicillin Streptomycin (Gibco, 15140-122), and 50 µM 2-mercaptoethanol (ThermoFisher, 31350010). The cells were fed every four days, and iMac precursors were collected for culture supernatants from week 2 to week 5. Differentiation of iMacs from precursors was achieved by culturing cells in RPMI-1640 media (Gibco, 21875-034), supplemented with 10% FBS (Sigma-Aldrich, F9665), 2 mM L-Glutamine (Sigma-Aldrich, G7513) and 100 U/mL Penicillin and 100 μg/mL streptomycin (Gibco, 15140-122), and human M-CSF at 5 ng/mL (Peprotech, 30025). Assessment of different M-CSF concentrations on PIEZO1 induced phenotypes were tested before selecting the optimal dose for experiments (Fig. S2D-E).

### Long-term and short term PIEZO1 activation with Yoda1

*Long term.* The iMacs precursors were isolated from the EB bodies as described above and cultured in complete RPMI medium with 5 ng/mL M-CSF for 72 hours. The cells were stimulated with Yoda1 at 15 μM for another three days before cell phenotype assessment by flow cytometry. Monocyte-derived macrophages were cultured in media as above supplemented with 5-20 ng/mL M-CSF for 3 days, and then stimulated with Yoda1 at 15 and 20 μM for another three days before flow cytometry assessment. In macrophage cultures, the media was replenished at day 3 with new M-CSF supplementation. In all experiments, cells cultured with equivalent to Yoda1 concentration of DMSO or medium were used as controls. *Short-term.* Macrophages were differentiated from CD14 monocytes as described above for 6 days in complete RPMI media supplemented with 5 or 20 ng/mL M-CSF. On day 5 of differentiation, the cells were stimulated with 20 μM Yoda1 for 16 hours followed by flow cytometry assessment. In some long-and short-term experiments, cells were also activated with 1ng/ml LPS (Sigma L6529, clone 055:B5) for 12 hours before assessment by flow cytometry.

### Resazurin Assay

To assess the cytotoxicity of PIEZO1 activation by different concentrations of Yoda1, we performed cytotoxicity assay using resazurin Alamar Blue HS assay (Thermofisher, A50100). Briefly, cells were seeded in 96 well plates at densities of 2×10^4 cells/well in 300uL of medium. In the wells containing the cells, 100 μL of fresh media with 10% AlamarBlue was added to the cells. The plate was incubated at 37℃ for 4 hours. Fluorescent intensity (FI) was measured for 560nm excitation and 590nm emission. The viability was calculated as percentage as per manufacturer’s instructions: (FI from Yoda1 cells/FI from DMSO controls) x100.

### Calcium Flux Assay

Day 3 macrophages were lifted using Accutase and seeded in a 96-well plate at a density of 4 x10^4^ cells per well. On Day 6, cells were stained with Fluo-4 NW Calcium Assay Kit (Invitrogen, F32605) as per manufacturer’s protocol. Briefly, assay buffer was mixed with probenecid stock to make dye loading solution, cells were washed twice with PBS, then 100µL dye loading solution was added to each well and incubated for 30 minutes at 37°C, followed by a further 30 minutes at room temperature. Dye loading solution was quickly removed before immediately adding Yoda1, PBS/DMSO or ionomycin using a multichannel pipette to the wells. The plate was analysed using a VANTAstar plate reader (BMG Labtech).

### BD Rhapsody Single cell transcriptomics of PIEZO1 activated macrophages

M-CSF (20 ng/ml) driven CD14 monocyte cultures were stimulated with 20 μ M Yoda1 at day 3 for 72 hours or at day 5 for 12h. The cells were lifted by incubation with Accutase (Sigma-Aldrich A6964) for 8 minutes at 37°C. The cell suspension was then transferred into the respective FACS tubes, cells were counted using a haemocytometer and 3000 cells from each donor at each condition was added to separate 1.5 mL tubes. The cells were then labelled by incubation with unique tags (BD Human multiplex tags, BD, 633781) for 20 minutes at RT. The cells were then washed three times, pooled together, and loaded onto the scRNA-seq BD Rhapsody Cartridge, using the BD Rhapsody Cartridge Reagent Kit (no. 633731) according to the manufacturer’s protocol. The cDNA was synthesised using BD Rhapsody cDNA Kit (Cat. No. 633773), the whole transcriptome and Sample Tag libraries were prepared using the whole transcriptome and sample tag library Preparation kit (BD Cat. No. 633801), as per manufacturer’s protocol (2019 version). Sequencing was performed using Illumina system sequencing (Glasgow Polyomics) Mapping was performed on the BD Velsera website, using the BD Rhapsody Sequence Analysis Pipeline, version 1.2. For whole transcriptome mapping with introns, FASTQ files containing the raw sequencing were used to map against an annotated reference genome (RhapRef_Human_WTA_2023-02.tar.gz). The sample Tag information can be found in **Table S1.** The pipeline generates a Seurat object as RDS file.

### Analysis of single-cell transcriptomics

Analysis of the data has been performed using the R version 4.2.2. The Seurat package v4 was used to analyse an object read (readRDS) from the RDS file from SevenBridges mapping. Cells of undetermined origin and multiplets were excluded using subset (subset, invert = True), based on annotation in the Seurat object from the mapping analysis. The data was normalized (NormalizeData) and the top variable genes were identified for all samples (FindVariableFeatures). The data were scaled (ScaleData) and principal component analysis was run (RunPCA). Metadata were added to the object as described in **Table S1-3.** *Quality control and filtering*. Cell filtering included removal of cells with the following expression levels of mRNA and mitochondrial genes (subset). For experiment 1: nFeature_RNA > 2200 & nFeature_RNA <6000 & nCount_RNA > 200 & nCount_RNA < 37000 & percent.mt < 20). Experiment 2: nFeature_RNA > 2200 & nFeature_RNA <6000 & nCount_RNA > 200 & nCount_RNA < 37000 & percent.mt < 17). Experiment 3: nFeature_RNA > 1900 & nFeature_RNA <5600 & nCount_RNA > 200 & nCount_RNA < 27000 & percent.mt < 17). Thresholds were determined by visualising using a violin plot (VlnPlot) of features such as nFeature_RNA, “nCount_RNA and “percent.mt”. This approach excluded doublets and dying cells. *Clustering pipeline.* The top 40 PCA embeddings were visualised and then used for Uniform Manifold Approximation and Projection (UMAP) generation (RunUMAP). The same PCs were used to determine the k-nearest neighbours for each cell during SNN graph construction before clustering at the chosen dimension and resolution (FindNeighbors, FindClusters). The objects were saved for integration. Clusters were visualised in UMAP graph (Dimplot).

#### Harmony integration

The three experiments RDS objects (readRDS) were merged into one (merge). The data were normalised (NormalizeData) and the top variable genes were identified for all samples (FindVariableFeatures, selection.method vst and nfeatures 2000). The row names of the object were assigned to a new variable called all.genes. The data were scaled (ScaleData, features = all.genes). Principal component analysis was run (RunPCA, npcs = 40). The merged object was Integrated using harmony Click or tap here to enter text.(Korsunsky et al., 2019)(RunHarmony) grouping by variable Donor and Experiment **(Table S1**), and theta =2 for both variables. *Clustering pipeline for integrated object.* The FindNeighbors function was applied to the integrated Seurat object, using the "harmony" reduction. The top 27 harmony embeddings were used to calculate the neighbour graph. Then the FindClusters function was run on the object, also using the "harmony" reduction and a resolution parameter of 0.5 was selected. The RunUMAP function was then executed on the object, leveraging the "harmony" reduction and the first 27 dimensions to compute an UMAP. *Differential Marker Analysis of the clusters*. Marker genes for each cluster in the integrated Seurat object were identified using the FindAllMarkers function. This function was configured to identify only positive markers (only.pos = TRUE) and include features expressed in at least 25% of cells (min.pct = 0.25). Significant markers were filtered using a log fold-change threshold of 0.25 (logfc.threshold = 0.25). The identified markers were further filtered to retain only those with an adjusted p-value less than 0.05. This step ensures that only statistically significant markers were included in the downstream analysis.

#### RNA Velocity of PIEZO1 activated macrophages

The Seurat object was converted to an AnnData object for downstream analysis in Python using the SeuratDisk package (v0.0.0.9020) (https://github.com/mojaveazure/seurat-disk). RNA velocity was inferred on the whole dataset by first estimating the spliced and unspliced counts using the ‘*run 10X’* command from the velocyto package (La Manno et al., 2018). Loom files were generated for each of the three experiments and were merged using the loompy package (v3.0.6) and subsequently merged with the single cell data using scVelo (v0.3.2) (Bergen et al., 2020) to create a separate layer for the spliced and unspliced counts. Data were next subset to include key macrophage clusters of interest. The two splice layers were then normalised and underwent the recommended preprocessing steps as described in the scVelo documentation before proceeding with the RNA velocity analysis using CellRank2 (v2.0.6) (Bergen et al., 2020). To then quantify cellular fate, transition matrices were computed with the VelocityKernel and RealTime kernel using a weight of 0.8 and 0.2 respectively. The RealTime kernel was initialised using the experimental timepoint values in the data to allow for a temporal axis in the combined kernel. Initial and terminal macrostates were then estimated using the GPCCA (Generalized Perron Cluster Cluster Analysis) estimator which identified distinct cell states. After the key macrostates were determined using an RNA-Realtime combined kernel, pseudotime was calculated using the Palantir (5) package following the recommended steps outlined in its documentation. Diffusion components were run for the first 5 components and data was imputed using MAGIC (van Dijk et al., 2018)for downstream visualisation of gene expression trends. A PseudotimeKernel was then constructed using the values computed by Palantir implementing the lineage states calculated from the experimental time-based RNA velocity analysis. Finally, putative lineage drivers of the constructed trajectory were identified, ordered by Palantir pseudotime values and visualised using the CellRank2 function (*pl.heatmap*).

### Bulk RNA sequencing

Macrophages were stimulated with Yoda1 or DMSO control for 12 hours as described above. Cells were lysed using using RLT buffer containing 1% Beta-mercaptoethanol. RNA was then extracted using the RNeasy Micro kit (Qiagen, 74004). RNA Libraries were prepared and sequenced by the MVLS Shared Research Facility at the University of Glasgow using Illumina NextSeq500. The data processed by the MVLS Shared Research Facility. In brief, the FASTQ files were quality control checked using FastQC (Andrews, GitHub, v0.12.1: https://github.com/s-andrews/FastQC), reads were then trimmed using FastP (Chen et al., 2018) and then aligned to the human reference genome (Ensembl, release 112) using HISAT2 (v2.2.1) (Kim et al., 2019). Read counts were generated using featureCounts (Liao et al., 2014). The normalised expression and differential expression values were generated using DESeq2 (v1.44.0) (Love et al., 2014)with default settings and no additional covariates. The data was analysed and visualised using RStudio and applying a significance threshold of adjusted p < 0.05 and absolute log2fold > 0.58. The GO biological processes database was used for over-representation analysis, at a significance threshold of adjusted p < 0.05.

### Flow cytometry analysis of macrophage phenotypes

The cells were washed with D-PBS, and detached cells were collected into individual FACS tubes (Falcon, 352052). The remaining adherent cells were det-attached by incubation with Accutase (Sigma-Adrich, A6964) at 37°C for 8 minutes. The cell suspension was then transferred into the respective tubes, and the wells were washed once with D-PBS to collect remaining cells. The cells were then labelled with the viability dye eFluor780 (Invitrogen, 65086514) in D-PBS in the dark on ice for 20 minutes. After the incubation period, cells were washed by adding 1mL of FACS buffer and centrifugation at 1600 RPM for 5 minutes at 4oC. FACS buffer contained 1X D-PBS with 2% FBS (Sigma-Aldrich, F9665), 2mM L-Glutamine (Sigma-Aldrich, G7513-100ML), 100 U/mL Penicillin and 100 μ g/mL streptomycin (Gibco, 15140-122). The supernatant was discarded, and cells were labelled with a mix of antibodies against markers of interest **(Table S6)** in the dark on ice for 30 minutes. Unlabelled cells, cells labelled only with viability dye and cells with FMO for each marker of interest were used as technical controls for gating. The data acquisition was performed using BD LSR FORTESSA cytometer. The results were analysed using FlowJo_v10.8.0.

### Evaluation of IL-8, CCL3 and TNF

Cell culture supernatants were centrifuged at 2000xG for 10 minutes to ensure no cells were present. After centrifugation, the supernatant was carefully collected into new sterile 1.5mL tubes, avoiding disturbance of the pellet. TNF (Thermofisher, CHC1753), IL-8 (Thermofisher, 88808688) and CCL3 (Thermofisher, 8873522) ELISA kits were used according to the manufacturer’s protocol.

### Immunofluorescent staining of cultured cells

CD14 monocytes were isolated as described above and were cultured in Ibidi 8-chamber slides at starting densities of 2×10^5 cells/well in complete RPMI 1640 with 50 ng/ml M-CSF for 6 days and stimulated as described in results. After removing media, cells were fixed in 4% methanol-free PFA at RT for 30 minutes. The cells were washed 4 times in D-PBS and then permeabilised by incubation with 0.025% Triton-X in PBS for 30 minutes. The cells were washed 3 times and blocked in 1% BSA in PBS with 10% human serum (Thermofisher, 31876) and 10% goat serum (Thermofisher 31872). After blocking, the slides were washed twice in PBS and incubated with primary antibodies or isotope controls in 1% BSA/PBS were (Supplemental Table 6) overnight at 4°C. The following day, the slides were incubated with the secondary antibodies for 2 hours at RT. The cells were then washed three times. Silicon chambers were removed and slides were mounted using VECTASHIELD DAPI mounting medium (2B Scientific, H-1800-10).

### Immunofluorescent and RNAscope staining of paraffin-embedded sections of synovial tissue biopsies of RA patients

#### Synovial biopsies

Patients fulfilling the EULAR classification criteria revised criteria for RA (Aletaha et al., 2010) and individual fulfilling the EULAR classification criteria for Clinical Suspect Arthralgia at risk of RA development (van Steenbergen et al., 2025) were enrolled and underwent ultrasound-guided synovial tissue biopsy as a part of ongoing recruitment to the SYNGem cohort (Alivernini et al., 2021) according to the standard-of-care protocol, Division of Rheumatology and Clinical Immunology, Fondazione Policlinico Universitario A. Gemelli IRCCS – Università Cattolica del Sacro Cuore, Rome - Italy. RA patients with active disease were treatment-naïve or in sustained clinical and ultrasound remission as previously defined (Alivernini et al., 2017). The clinical and laboratory evaluation of each RA patient enrolled included DAS based on the number of tender and swollen joints of 44 or 28 examined, plus the erythrocyte sedimentation rate (ESR) and plasma C-reactive protein (CRP). Peripheral blood samples were tested for IgA-RF and IgM-RF (Orgentec Diagnostika, Bouty-UK), and ACPA (Menarini Diagnostics-Italy) using commercial Enzyme-Linked Immunosorbent Assay (ELISA) and ChemiLuminescence Immunoassay (CLIA) respectively. Demographic, clinical, and immunological features of this cohort are shown in **Table S7**. In addition, synovial tissue was obtained from individuals undergoing arthroscopy for meniscal tears or cruciate ligament damage with macroscopically and MRI-confirmed normal (“healthy”) synovium (**controls, n = 2**) at the Division of Sport Traumatology and Joint surgery, Fondazione Policlinico Universitario Agostino Gemelli IRCSS, Rome, Italy. The demographic characteristics of this cohort are presented in Supplemental Table 7. The study protocols on adult RA patients’ and healthy donors’ synovium were approved by the Ethics Committee of the Università Cattolica del Sacro Cuore (14996/20 and 00243/23). All samples were taken with written and informed consent obtained from all sample donors.

#### Immunofluorescence staining

Formalin-fixed paraffin-embedded 5 μ m-thick synovial tissue sections deparaffinized and dehydrated. Antigen retrieval was performed using 0.01M citrate buffer with pH 6.0. Blocking was performed with TBS containing 10% human serum, 10% goat serum and 1 % BSA. After blocking, slides were stained with primary antibodies PIEZO1 (Biolegend, 602752) and CD68 (Dako, M0876) or isotype (Invitrogen, 14-4714-82) at 4°C overnight. The following day, slides were incubated with the secondary antibodies for 2 hours at RT. Nuclei staining was performed using DAPI mounting medium and slides were analysed using Confocal microscopy (Zeiss LSM880).

#### RNA scope staining

For detection of PIEZO1 transcripts in synovial tissue macrophages, we used the RNAscope Multiplex Fluorescent v2 assay (Advanced Cell Diagnostics, Newark, CA, USA; cat. no. 323100) on formalin-fixed paraffin-embedded (FFPE) synovial tissue sections. Tissue blocks were sectioned at 4–5 μm, deparaffinized, and processed for target retrieval and protease treatment according to the manufacturer’s instructions. The PIEZO1 probe (cat. no. 485101) was hybridized to the sections and visualized using the C2 fluorescent channel (AF647, 1:5,000 dilution). Following in situ hybridization, sections were immune-stained with Alexa Fluor® 594 anti-CD68 antibody (clone KP1 at 1:100 dilution; cat. no. ab2772938, Abcam) to identify macrophages. Nuclei were counterstained with SYTO™ 83 Orange Fluorescent Nucleic Acid Stain (1:1000 dilution) (ThermoFisher, S11364). Slides were analysed using GeoMX Digital Spatial Profiler (Nanostring).

##### GeoMX Digital Spatial Profiling

Formalin-fixed, paraffin-embedded (FFPE) tissue blocks were sectioned at 5 μm thickness onto Superfrost Plus microscope slides. Sections from two patient samples were mounted on the same slide, depending on tissue size. Slides were processed for DSP tissue treatment within 1 hour of sectioning to preserve RNA quality. For NanoString GeoMx DSP RNA assays, slides were prepared according to the FFPE RNA Slide Preparation Protocol described in the GeoMx NGS Slide Preparation User Manual (NanoString, MAN-10150-07). Briefly, following target retrieval (15 minutes in Tris-EDTA at 99°C), slides were incubated in Proteinase K (1 μg/ml for 15 minutes) and then hybridized with the GeoMx Cancer Transcriptome Atlas Panel (1811 targets; cat. 121400101). Each slide was covered with 200 μL of prepared probe hybridization solution, overlaid with a HybriSlip (Grace Bio-Labs, cat. 714022), and incubated overnight at 37 °C. Following hybridization, HybriSlips were removed by immersing slides in 2× saline-sodium citrate (SSC; Sigma-Aldrich, cat. S6639). To remove unbound probes, slides were washed twice in stringent wash buffer (50% formamide [Thermo Fisher, cat. AM9342] in 2× SSC) at 37 °C for 25 min.

Slides were then blocked with 200 μL of Buffer W (NanoString) in a humid chamber for 30 min at room temperature. Following in situ hybridization, sections were immunostained with Alexa Fluor® 594 anti-CD68 antibody (clone KP1; cat. no. ab282650, Abcam) to identify macrophages. Nuclei were counterstained with SYTO™ 83 Orange Fluorescent Nucleic Acid Stain (ThermoFisher, S11364). Regions of interest (ROIs) were selected based on morphology and marker staining (e.g., lining and sublining CD68⁺ cells). Libraries were prepared using MAN 10153-08 and sequenced using NovaSeq 6000 (Illumina) instrument.

### Intraarticular activation of Piezo1 in healthy mice

To assess the activation of PIEZO1 on the synovial tissue macrophages, 5 μM Yoda1 in D-PBS was injected intraarticularly in WT C57BL/6 mice in one knee, whereas the vehicle (0.025% DMSO in D-PBS) was injected in the opposite knee. Two injections were performed in each knee three days apart. The synovium was extracted from each knee of mice 24 hours and 72 hours after the second injection.

### Collagen Antibody Induced Arthritis (CAIA) model

The mice were injected Arthritomab cocktail (mdbioproducts, CIA-MAB-2C) intravenously (4mg/mouse) on day 0. On day 2 after antibody injection, the mice were injected intraarticularly into contralateral joints with Yoda1(5μM) or vehicle control as described above. On day 3, 100 μg/mouse LPS was injected intraperitoneally, to disease onset. On day 4, the mice received a second dose of Yoda1(5μM) or vehicle. Arthritis scores were recorded each day. The synovium was extracted from each knee 24 hours later (day 5=peak of CAIA inflammation, N=6 mice) and 72 hours later (day 7, N=6 mice) after the second Yoda/Vehicle injection.

### Digestion of mouse synovium

Mouse synovia were collected into 20ml universal tubes with serum-free RPMI 1640. The synovia were then transferred in fresh universal tubes containing 4.8 mL of serum free RPMI 1640 and supplemented with 150 µL Liberase TM (Merck, 5401119001) (working concentration 150 µg/mL), and 42 µL DNase I (Qiagen, 79254) (working concentration 11 µg/mL). Tissues were incubated on shaker at 195 rpm at 37 ℃ for 35 minutes. After digestion 5 mL of cold complete RPMI 1640 containing 10% FBS was added to the cells to terminate enzymatic digestion. Using 100 µm filter, the cells were passed into a new 50 mL tube. Using a small syringe plunger, the remaining tissue were mechanically pressed against the filter, to release extra cells into a sterile petri dish. The extra cells were collected in 1 mL of complete medium and added to the cell suspension. The cell suspension was spun at 420 g at 4 ℃ for 8 minutes. The cells were then resuspended in FACS buffer for staining with specific antibodies **(Table S6)** as described above.

### Mouse joints collection and decalcification for histology

After excessive muscle removal, the mouse legs were fixed by submerging in 4% PFA for 24 hours. The PFA was removed, and legs were stored in 100% ethanol until ready for decalcification. To start the decalcification, the legs were submerged in 10% EDTA with pH 7.4. The legs were decalcified in EDTA solution in a shaker, at 4°C. The EDTA was changed every 3 days, until the legs became elastic (approximately 10 days). The decalcified legs were dehydrated using an automatic ethanol gradient processor, incubated in Xylene and embedded in paraffin blocks. Joints embedded in paraffin wax were cut in sagittal sections (6 μm), then stained with haematoxylin, safranin-O/fast green. The slides were scanned and assessed using Zeiss Zen software.

### F-actin staining of macrophages

#### Monocyte-derived Macrophages

Monocytes were isolated from PBMCs and seeded to ibidi 8-chamber slide at a starting density of 1.5×10^5 cells/well. Monocytes were differentiated to macrophages for 6 days with 20 ng/mL M-CSF. The cells were then stimulated with Yoda1 at concentration of 20 μM for 1, 5, 15, and 30 minutes. At the end of each incubation period, they were fixed using methanol-free PFA, added to the media at a final concertation of 4% PFA for 15 minutes. After fixation, the cells were washed 3 times with D-PBS, and permeabilised with 0.02 % Triton-X in D-PBS for 10 minutes at RT. The cells were washed three times with PBS and then stained using phalloidin ActinGreen Probes in D-PBS (Thermofisher, R37110), for 30 minutes in the dark as per manufacturers instruction. The cells were then washed three times. Silicon chambers were removed and slides were mounted using VECTASHIELD DAPI mounting medium (2B Scientific, H-1800-10). Images were taken using confocal microscope and F-actin intensity was analysed as described above.

### Calpain inhibition assay

CD14 monocytes were cultured for 6 days in complete RPMI with 20 ng/mL M-CSF. For calpain inhibition the cells were pre-incubated with 100 µM Calpain inhibitor PD150606 (Sigma, 513022) for 2h. After that 20µM Yoda1 was added to the cells. After 12 hours, the media containing calpain inhibitor and Yoda1 was removed, and cells were cultured in complete RPMI with M-CSF for another 12 hours. The cell phenotypes were assessed by flow cytometry as described earlier.

### Western Blot for HIF1A

Macrophages were differentiated from CD14 monocytes as described earlier. On day five, the cells were incubated with PBS/DMSO or 20 µM Yoda1 for 12 hours. Following stimulation, the cells were lysed by incubation in complete RIPA lysis buffer (0.25% Sodium Deoxycholate, 50 mM Tris-HCl pH 7.4, 1% IGEPAL, 150 mM Sodium Chloride, and 1 mM EDTA pH 8.0) containing protease inhibitors (1 mM Sodium Fluoride, 1 mM Sodium Orthovanadate, 1 mM PMSF, 10 µM Leupeptin, 1 µM Pepstatin A, and 0.3 µM Aprotinin) for 30 min on ice. The concentration of protein samples was quantitated using Bradford protein assay (ThermoFisher, 23200). Denatured protein samples were loaded onto the SDS-PAGE gels (5% stacking gel and 10% resolving gel) for electrophoresis. Separated proteins were then transferred onto 0.2 µm nitrocellulose membrane (Cytiva, 10600001) at 140 V for one hour, blocked with 5% skim milk in PBS-Tween 20 buffer. Primary antibodies, HIF1A (CAT#14179, CST) and β - actin (GenScript, A00702) were probed. Blots were imaged using Li-Cor Odyssey DLx and c-Digit models where appropriate.

### CRISPR-Cas9 mediated knock-out of HIF1A in CD14 monocytes

The guides (a premix of three guides: AGGAAAGUCUUGCUAUCUAA, ACACAGGUAUUGCACUGCAC, and UUCACAAAUCAGCACCAAGC) purchased from Edico, were mixed with the Cas9 (Cas9 / SpCas9 2NLS Nuclease) at a molar ratio of 5:1 in an RNase-free low-bind tube and incubated at room temperature for 30 minutes for the development of ribonucleoprotein complex (RNP). Lonza P3 buffer was prepared by mixing P3 buffer with the supplement as follows: 82 µL Lonza solution and 18 µL supplement for a total of 100 µL suspension, and kept at RT. The buffer was then mixed with the RNP complexes at the end of the 30 minutes incubation. This mix was then used to resuspend the cell pellet, and the suspension was immediately transferred to a 100 µl Lonza cuvette. Cells were electroporated using the CM137 pulse code, in Amaxa 3D Nucleofector. After the successful electroporation, the cells were immediately transferred to pre-warmed complete media containing 20 ng/mL M-CSF and cultured for 6 days. To validate the knockouts, the genomic DNA was extracted (Invitrogen, K182001) from WT and HIF1a-KO macrophages after 3 days in culture with 20 ng/mL M-CSF. The PCR was performed using DreamTaq PCR mastermix (Thermofisher, K1081) with HIF1a forward primer (TCAAAAACGTGATTAATCCACTAGTGA) and reverse primer (GTGTGAGCCACCATGCAT).

The PCR products were purified using Monarch clean-up kit as per manufacturer’s protocol (Monarch, T1030S). A mix of the purified PCR product with sequencing primer (CGTGATTAATCCACTAGTGACAGTAAATTT) was prepared using the Mix2seq kit and sequenced by Eurofins. Sequencing .abi files from KO and WT cells were uploaded to ICE website (Edico) and population wide KO was confirmed, showing multiple indels in the KO compared to the WT sequence.

### Evaluation of the expression of markers on paraffine sections and slides

#### Quantification of PIEZO1 in synovial tissue from RA patients

The images were analysed using ImageJ with plugin Bio-formats, allowing upload of .czi files and slitting of fluorescence channels. Starting from cells stained with macrophage marker CD68, regions of interest (ROIs) were drawn around macrophages and saved. Then these ROIs were mapped into the respective PIEZO1 channel, from the same image. The Raw Integrated Density and Area were measured in the PIEZO1 channel, using ImageJ Measure for each ROI. In addition, 10 random background measurements were performed from the areas outside of the tissues.

The corrected total cell fluorescence (CTCF) was calculated using the formula: CTCF Integrated Density – (Area of selected cell X Mean fluorescence of background readings), as described in literature (Burgess et al., 2010; McCloy et al., 2014). This gives a corrected intensity value for PIEZO1 expression in each macrophage. In the end, the mean of CTCF from multiple macrophages in the lining layer was analysed per each donor (n=5), and compared between At-risk, Active disease and Remission.

#### Evaluation of TJP1 in STM-resembling iMacs

Cells were fixed and stained for TJP1 and F-actin. Using Imagej with Bio-formats as above, the channels were split and ROIs were generated based on the F-actin channel, to cover the whole shape of the cell. The ROIs were added into the TJP1 channel and Area and Raw Integrated Intesity were measured. Background measurements were collected randomly from areas outside of cells. The CTCF was calculated as described above.

#### Quantification of F-actin in cultured macrophage stimulated with 20μM Yoda1 at different timepoints

F-actin was quantified using ImageJ per total field of view in images from 3 experimental repeats (n=3) for each timepoint. F-actin was quantified as follows: Intensity= RawIntegratedDensity /(Area/number of cells), using ImageJ software. Number of cells was determined by counting the nuclei per each image/total field of view.

#### Calculation of nuclear circularity and size

Nuclear shape and size were calculated in macrophages from F-actin experiment above. The nuclei were identified by positive staining of DAPI, when splitting the channels using ImageJ Bio-formats, and ROIs were generated around their perimeter. A total of 45 random nuclei was chosen for each condition (e.g. Yoda1/DMSO) and timepoint. Area and Perimeter were measured using Image J. Circularity was calculated using the formula 4π(area/perimeter^2).

#### Measurement of macrophage spindle circularity

Macrophage spindle circularity was measured from cultures of cells from five donors stimulated with 20µM Yoda1 or 0.1% DMSO for 72 hours. The light microscopy images were pre-processed using Gaussian Blur and thresholding to enhance spindle cell visibility, followed by binary conversion and watershed segmentation to separate adjacent cells. Cell counts were obtained using the Analyze Particles function in ImageJ, applying size and circularity filters (circularity: 0.1–0.5) needed for assessment of the elongated morphology of spindle cells.

### Ethical approval

All patients with RA fulfilled the 2010 American College of Rheumatology (ACR)/European Alliance of Associations for Rheumatology criteria for RA (Aletaha et al., 2010) while individuals at risk of RA met EULAR/ACR risk stratification criteria for development of rheumatoid arthritis in the risk stage of arthralgia (van Steenbergen et al., 2025). Patients were recruited at Division of Rheumatology and Clinical Immunology, Fondazione Policlinico Universitario A. Gemelli IRCCS – Università Cattolica del Sacro Cuore, Rome under the ethical approval of the Comitato Etico Territoriale Lazio Area 3 (14996/20 and 00243/23). Demographic and clinical characteristics of the RA patients are shown in **Tables S7.** Healthy controls were recruited at the Division of Sport Traumatology and joint surgery, Fondazione Policlinico Universitario A. Gemelli IRCCS, Rome, Italy, under ethical approval the Comitato Etico Territoriale Lazio Area 3 (00243/23). Blood from healthy controls were collected at the University of Glasgow by a trained phlebotomist, under ethical approval “Cells and Soluble Mediators in Peripheral Blood of Normal Healthy Donors, V6.0. (Study protocol on blood cells was approved by University of Glasgow, School of Infection and Immunity Ethical Committee (CG_2022_20_A).

### Statistical analysis

Detailed statistical methods are provided in each figure legend and in the scRNAseq method sections above. For single-cell RNA analysis, the default Wilcoxon rank-sum test was used to calculate the DEGs (Seurat, FindAllMarkers), and p-adjusted values of <0.05 were considered significant. For Bulk RNAseq, DESeq2 test was used to calculate the p-adjusted values. Mixed Linear Model was used to analyse GeoMX data. All data were analysed for normality using the Shapiro-Wilk test.

## Data Availability

All raw data are available to readers in ArrayExpress under accession number **E-MTAB-15203** (bulk RNAseq of PIEZO1 activated macrophages) **and E-MTAB-15200** (scRNASeq data of PIEZO1 activated macrophages.

## Funding

This work was supported by

The British Society for Immunology Career enhancing Grant to T.S

The TRAM Fellowship Grant (KENN no. 20 21 05) to M.D

Versus Arthritis UK grant no. 23229 to M.K-S and S.A

FOREUM grant no. 063 to M.K-S and S.A

FOREUM grant no. 073 to S.A and M.K-S

We thank University of Glasgow Shared Research Facilities and Clinical Research Facility for their support and Andrew McGucken for some of the blood samples.

## Author contributions

**TS:** initiated study concept and manuscript writing; established experimental procedures for all *in vitro* cultures and single cell RNA sequencing (Fig. 1B, D, E, F, G, H; Fig. 2B-D; Fig. 4B-E; Fig. S1A-F; Fig. S2B, C, E); contributed to mouse experiments (Fig. 4A-V) and analysis, including synovial digestion and gating strategy (Fig. S3A-B); optimisation, performance and quantification of soluble mediators analysis (Fig. 2F, G); optimisation and performance of CRISPR-Cas9 methodology (Fig. 5L; Fig. S3E); performed histological staining and analysis of healthy and RA human synovium (Fig. 3C & E); secured funding for mouse and scRNA experiment; contributed to manuscript discussions and editing, including generation of all supplementary material.

**MD:** contributed to study concept and manuscript writing; contributed to generation of experimental procedures for *in vitro* cultures (Fig. 1 B, J; Fig. 2B, C, D; Fig. 4L; Fig. S1 A-C; Fig. S2 B, C, D, E); contributed to mouse experiments (Fig. 4A-L) and analysis, including synovial digestion and gating strategy (Fig. S3A,B); optimisation, performance and quantification of soluble mediators analysis (Fig. 2F & G; Fig. S2); contributed to generation of RNAscope images and analysis (Fig. 1A; Fig 3B & D); optimised and performed calcium flux and calpain inhibitor experimental methodology in macrophages (Fig. 1C, 4F & G); contributed to manuscript discussions and editing, including generation of supplementary material.

**CH:** Conceptualised and assisted in mouse experiments (Fig. 2A-L) and provided methodology for extraction of synovial tissue from mouse joints; contributed to histological analysis of murine joints (Fig. 2C); contributed to manuscript writing and editing.

**KW:** Assisted in acquisition and analysis of bulk RNAseq experiments in Yoda1 stimulated macrophages (Fig. 2A, E; Fig. 5E, H; Fig. S2A; Fig. S3D); generated and analysed bulk RNAseq data (Supplemental Table 4); generated pathway analysis of bulk RNAseq (Fig. S1G); conceptualised and contributed to design of Yoda1 DEG signature analysis (Fig. 1J); contributed to manuscript writing and editing.

**DS:** Performed integration of Yoda1 DEG signature score into published dataset (Fig. 1K, L); contributed to manuscript writing and editing.

**WHN & SP:** Assisted in Yoda1-macrophage monoculture experimental design and analysis (Fig. 2B); performance and quantification of soluble mediators’ analysis (Fig. 2F, G).

**KB:** Assisted with intra-articular injections and experiments of (Fig. 4); contributed to manuscript writing.

**LY:** Contributed to the *in vivo* experiment (Fig. 4).

**BDZ:** Optimised, performed and analysed Western Blot of Hif-1α in Yoda1-macrophage monoculture (Fig. 5I); contributed to manuscript writing and editing.

**CDM, DC:** Performed and analysed GeoMX data (Fig.5L), and RNAscope data (Fig.1A and Fig.3B and D)

**AF:** Optimised and performed MiloR analysis of Yoda1 macrophage culture (Fig. 1F); contributed to data visualisation; contributed to manuscript writing and editing.

**JF:** Analysed nature medicine STM clusters for expression of PIEZO1 (Fig. 3A); validated knock-out of Hif-1α in macrophages (Fig. 4L); contributed to manuscript editing and writing.

**LC:** Evaluated nature medicine cluster expression in active RA vs healthy or remission conditions using miloR (Fig. 3A); provision of biopsy resource and patient data; contributed to manuscript writing and editing.

**OH:** Optimised and performed Velocity analysis of Yoda1-macrophage monoculture (Fig.5J); contributed to manuscript writing and editing.

**LD:** Contributed to the isolation of synovial membrane from mouse joints (Fig.4)

**MRG, VAP, LP and IDM:** Collected synovial tissue biopsies from healthy, at risk of RA individuals as well as RA patients included in the study (Fig1A, Fig.2B-E and Fig.5L); followed up the RA patients in the clinic and performed clinical assessment for condition annotations (active and remission).

**SAt:** Facilitated use of the servers and hardware

**YY:** Contributed to manuscript editing and writing

**MAD:** Provided clinical setting

**TDO:** Facilitated the use of servers and hardware for omic data analysis

**JC:** Funded the bulk RNAseq experiments and sequencing;

**RJC:** Supervised DBZ and generation of CRISPR Cas9 protocols; funded western blots

**CG:** Supervised AF, KW, CH, and YY; contributed to manuscript discussions

**LM:** Visualised PIEZO1 expression in TREM2+ macrophages from unpublished dataset; contributed to manuscript discussions, writing and editing.

**NB:** Supervision of MD; contributed to manuscript discussions, writing and editing.

**SA & MKS:** Initiated study concept, designed and supervised overall work, and wrote the manuscript.

---

**Figure S1.**
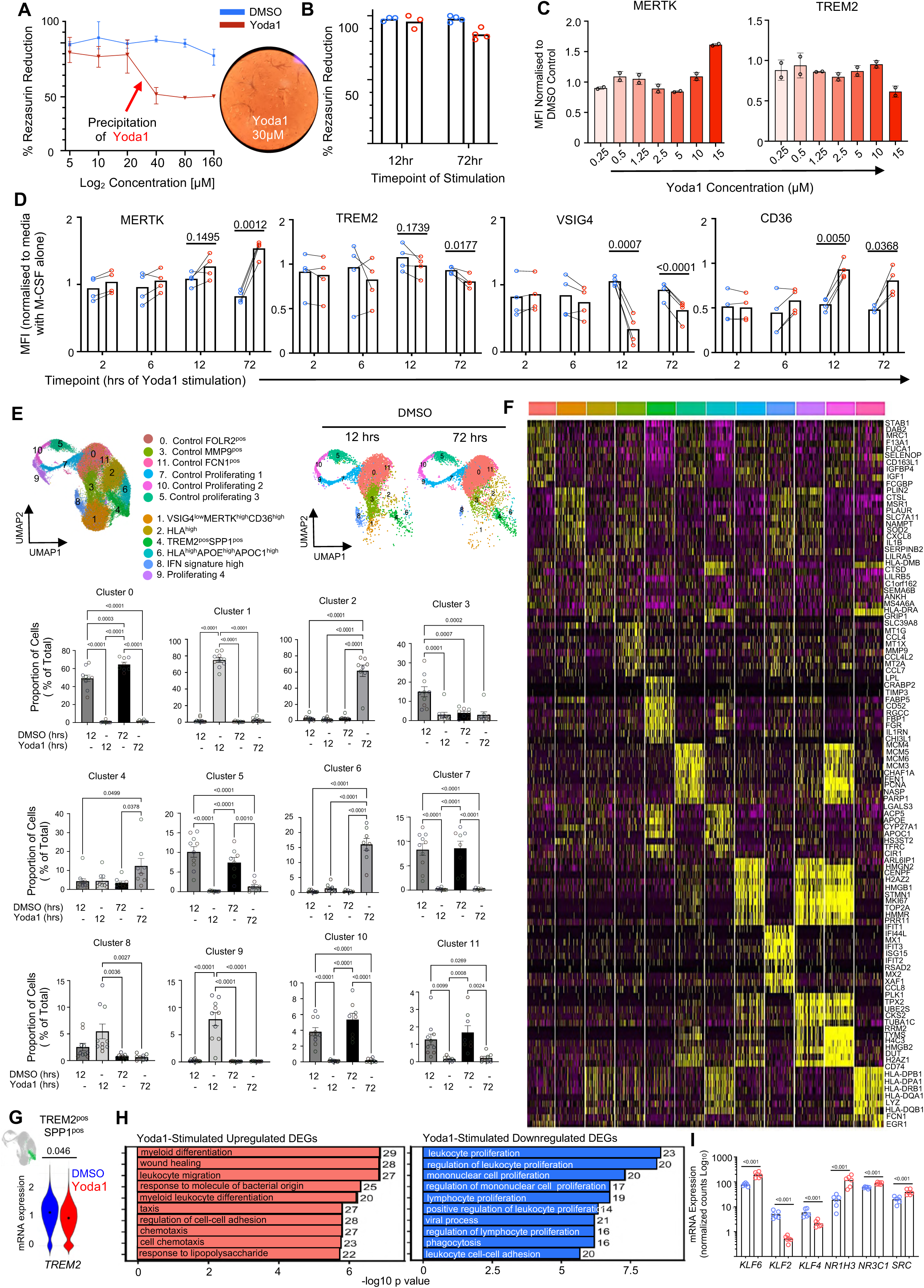
Characterization of monocyte- and iPSC-derived macrophage responses to PIEZO1 stimulation. **(A)** AlamarBlue Assay measuring % of Resazurin reduction in macrophages (n=3 donors) cultured with different concentrations of Yoda1 (5 – 160 μM) versus the equivalent DMSO controls (0.025- 0.8%) for 72h. Cytotoxicity was observed at concentrations of Yoda1 higher than 20µM. Precipitation of Yoda occurred at concentration of 30 μM for 72 hours. **(B)** Assessment of viability of monocyte derived macrophages (MDM, derived with 20ng/ml M-CSF) stimulated with Yoda1 (20µM ) for 12 and 72 hours. The short-term (12 hours) stimulation did not result in decreased viability of macrophages, and only 10% cell death was observed for longer term stimulation (72 hours), Data from n=3 Donors. **(C)** Assessment of the impact of Yoda1 (0.25 - 15 μM) on the expression of tissue-resident macrophage markers MERTK and TREM2 by flow cytometry. The MDM as in B were stimulated with the different Yoda1 concentrations, or equivalent concentration of DMSO for 72 hours. Values represent the MFI of these markers in CD11B^pos^ cells from N=2 donors. **(D)** Assessment of the impact of Yoda1 (20μM for 2, 6, 12 and 72 hours) on the mRNA expression of tissue-resident macrophage markers, MERTK, TREM2, VSIG4 and CD36 in monocyte-derived macrophages. Data shown paired values from N=4 donors, paired T test, the exact p value are on the graph. **(E)** UMAP visualisation of single cell transcriptomics of macrophages as in Figure 1. To the right, the UMAP split.by Condition (DMSO or Yoda1), below the proportion of distinct clusters, as percentage of all cells, split by stimulation duration. Each dot represents cells from one donor (N=8). One-way Anova with Tukey’s correction for multiple comparison. Exact p values are provided on the graphs. **(F)** Heatmap of top 10 upregulated genes per each cluster depicted in D. All upregulated differentially expressed genes were obtained using FindAllMarkers, with a logFC threshold of 0.25. For the heatmap, top10 genes were selected based on the highest expression in each cluster. **(G)** mRNA quantification of *TREM2* expression in TREM2^pos^SPP1^pos^ cluster between conditions (Yoda1 versus DMSO stimuation). Additional information in Table S3. Paired T test, with exact p value on the graph. **(H)** ORA pathway analysis of Yoda1 upregulated (n= 301) and downregulated (n= 199) genes during 12h stimulation shared between scRNAseq and bulkRNAseq approaches as in Fig1J. **(I)** mRNA expression of transcription factors regulated by PIEZO1 activation in macrophages stimulated with Yoda1 (20μM) for 12 hours. Data are representative of N=6 donors. mRNA was normalised to total counts. Data obtained by Bulk RNA sequencing; statistical significance determined using DEseq2 with adjustment for multiple comparison. P-value (<0.0001) are shown on the graph. KLF6=Krüppel-like factor 6, KLF2= Krüppel-like factor 2, KLF4= Krüppel-like factor 4, NR1H3=Nuclear Receptor Subfamily 1 Group H Member 3 (LXRa), NR3C1=Glucocorticoid Receptor.

**Figure S2:**
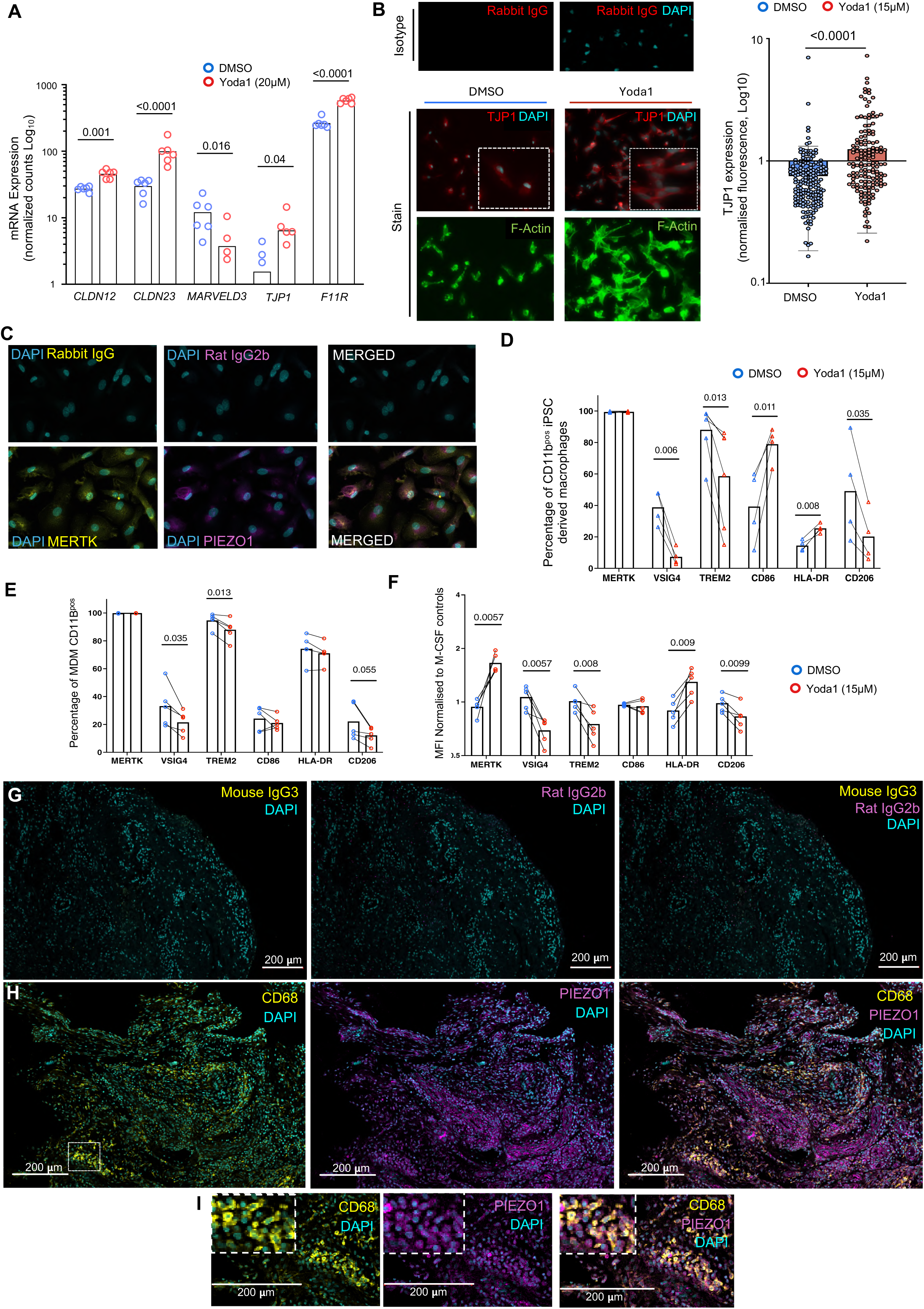
Expression of PIEZO1 and its downstream pathways in macrophages and in synovial tissue biopsies. **(A)** mRNA expression (bulk RNAseq) of tight junction proteins in macrophages stimulated with Yoda1 (20μM ) for 12 hours. Data are representative of N=6 donors. mRNA was normalised to total counts; statistical significance considered only for p-adjusted value < 0.05 (DEseq.). CLDN12=Claudin12, CLDN23= Claudin23, MARVELD3=MARVEL Domain Containing 3, TJP1=Tight Junction Protein 1, F11R=Junctional adhesion molecule A. **(B)** Quantification of TJP1 expression in iPSC-derived macrophages (iMacs) stimulated with DMSO (control) or Yoda1 (15µM). The cell borders were determined by staining of F-actin using AF488-conjugated phalloidin probes, and the corrected total cell fluorescence of TJP1 was calculated as described in previous work (doi: 10.4161/cc.28401). Images are representative of iMacs staining for TJP1 for Yoda1 and control (DMSO), in the two different experiments. Statistical analysis performed using unpaired T test. **(C)** Optimalisation of MERTK and PIEZO1 staining with specific antibodies in monocyte derived macrophages (20 ng/mL M-CSF for 7 days). Images are representative of macrophages from N=2 donors. Images taken with confocal microscope; images displayed at ‘maximum projection intensity’ using Zeiss Zen Black software. **(D)** Percentage of cells expressing STM markers in CD11B^pos^ iPSC-derived macrophages stimulated with 15µM Yoda1 or DMSO. Values are representative of N=5 experiments, normalized to control of cells in media with M-CSF (5 ng/ml); statistical analysis performed using paired T test. **(E)** Percentage of cells expressing STM markers in CD11B^pos^ monocyte-derived macrophages (MDM) stimulated with 15µM Yoda1 or DMSO. Values are representative of N=5 experiments, normalized to control of cells in media with M-CSF (5 ng/ml); statistical analysis performed using paired T test. **(F)** Median Fluorescent Intensity (MFI) of STM markers in monocyte-derived macrophages (MDM) stimulated with 15µM Yoda1 or DMSO. Values are representative of N=5 experiments, normalized to MFI M-CSF (5 ng/ml) controls ; statistical analysis performed using paired T test. **(G)** Optimalisation of PIEZO1 protein staining in human synovial tissue biopsies. Representative staining of macrophages with anti-CD68 (clone PG-M1), or its isotype Mouse IgG3 and anti-PIEZO1 (clone W17241A) and its rat IgG2b isotype. Images taken as tile scan at 20X, using confocal microscope and analyzed using Zeiss Zen Black software. **(H)** Representative **s**taining of CD68 (macrophages) and PIEZO1 in human synovial tissue from patients with active RA. Images taken as tile scan at 20X, using confocal microscope and analyzed using Zeiss Zen Black software. **(I)** Zoomed in image of single channels (Piezo1 and CD68) to show localization of the two markers in the macrophages.

**Figure S3.**
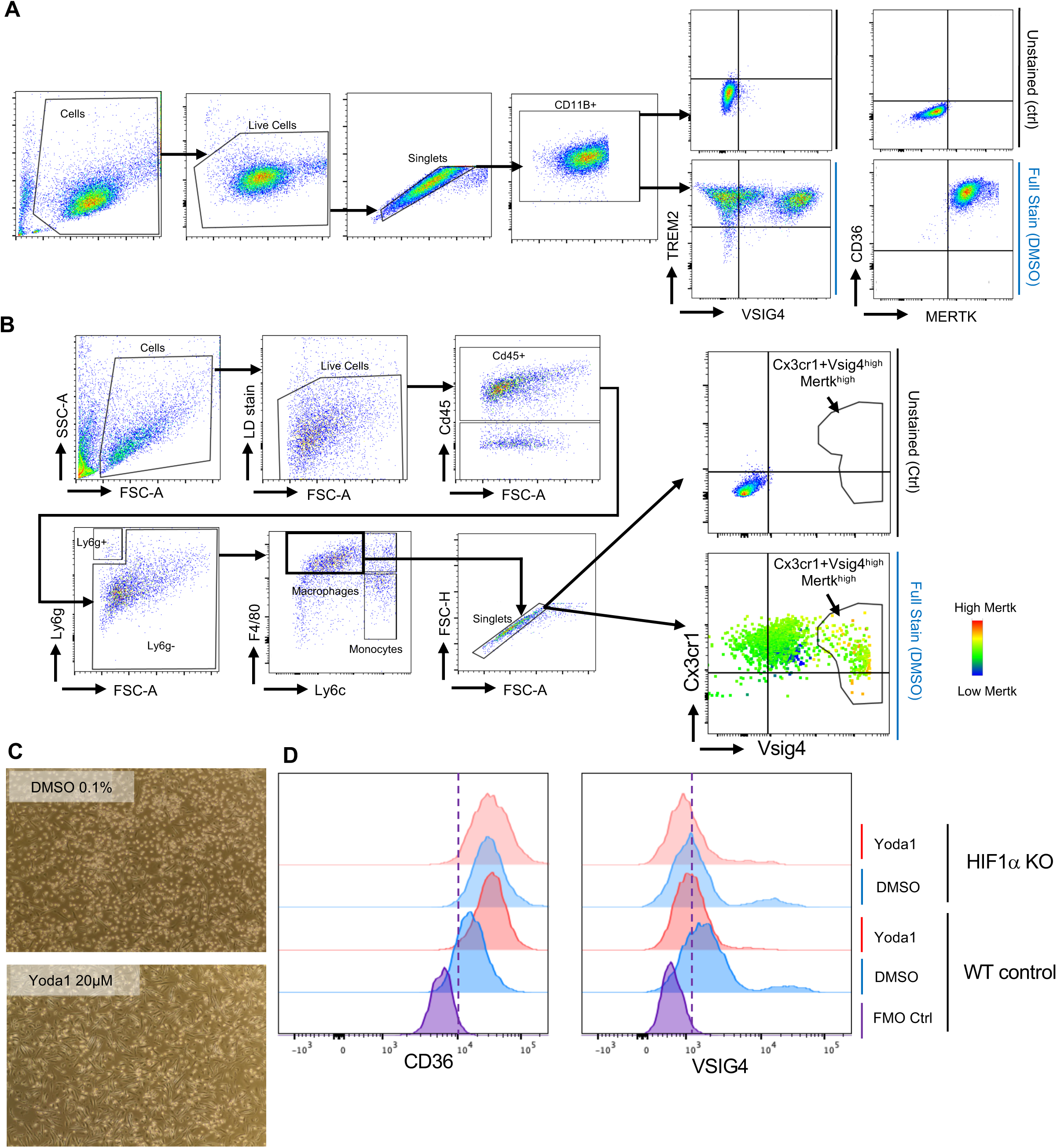
Gating strategy for mouse and human macrophages. **(A)** Gating strategy of Yoda1 stimulated human macrophages. Cells were gated based on side and forward scatter, then live cells were selected based on the lack of LD Efluor780 stain. Single cells were selected using a FSC area vs FSC height. Within the single cells, the CD11B positive cells were selected to gate the fully differentiated macrophages. Every marker of interest was gated within CD11B^+^ macrophages. Marker expression per cell was measured by the median fluorescent intensity (MFI) within the selected populations gate. FSC Forward scatter, SSC Side scatter, LD Live/Dead stain. **(B)** Gating strategy of distinct mouse synovial tissue cell population. Cells were gated based on side and forward scatter, then live cells were selected based on the lack of LD Efluor780 stain. In the Live population, haematopoietic cells were separated from stromal based on CD45 expression. In the Cd45+ population, neutrophils were identified by expression of Ly6g. Within the Ly6g negative population macrophage and monocyte were gated based on the expression of F4/80 and Ly6c, respectively. Macrophages were identified as F4/80+Ly6c-. Within this population lining-layer macrophages were identified based by Cx3cR1 and Vsig4 expression. **(C)** Light microscopy of MDM treated with 20 μM Yoda1 or 0.1% DMSO for 72 hours. The images are representative from one donor out of a total of N=5 donors. The quantification of cell spindle shape is shown in Fig 4. **(D)** Representative histograms of CD36 and VSIG4 expression in control and CRISPR-Cas9 guided HIF1α deficient macrophages. The histograms are representative of one donor out of a total of N=3 donor (Fig.5L)

